# Versatile and Scalable Reflective Micromirrors for Single-Objective Light Sheet Microscopy

**DOI:** 10.64898/2026.04.08.717282

**Authors:** Nahima Saliba, Siyang Cheng, Prakash Joshi, Anna-Karin Gustavsson

## Abstract

We present a tunable microfabrication pipeline for creating robust, reflective inserts that adapt conventional commercial imaging chambers for single-objective light sheet (LS) illumination. This system reduces the complexity associated with dual-objective LS setups and specialized LS chambers while retaining the native functionality and biocompatibility of the original chambers. The fabricated insert features a metalized, 3D nanoprinted micromirror with an angled reflective surface, enabling alignment of a thin LS for sectioning and imaging throughout mammalian cells. Using this pipeline, we demonstrate that single-objective LS illumination achieves an over 4X improvement in the signal-to-background ratio compared with conventional widefield epi-illumination in both fixed and live cell samples. Furthermore, we show substantial resolution enhancement for single-molecule localization microscopy compared to epi-illumination for improved imaging at the nanoscale. The versatile and scalable design offers an easily implemented approach to bring the benefits of single-objective LS microscopy to a wide array of biological studies.

## TEXT

Light sheet (LS) illumination, a technique that optically sections thick samples with a thin sheet of light, has proven to be a powerful tool for imaging challenging biological specimens by reducing fluorescence background, photobleaching, and photodamage^1–6^. Conventional LS setups typically use an illumination objective to generate the LS, which is positioned orthogonally to the focal plane of a separate detection objective^7–9^. While numerous modifications have been implemented to enhance the performance of the dual-objective design, such as deflecting the LS^10–13^, tilting the LS^14–16^, or employing a structured LS to form a lattice^17,18^, these configurations still present several challenges, including complex alignments, bulky setups, restrictive sample mounting requirements, and the risk of relative drift between the illumination and detection objectives during imaging. Furthermore, for live-cell imaging, these systems often necessitate intricate chambers to enclose the entire two-objective setup^19^ or constrained geometries where objectives are required to illuminate and detect through microfluidic chips or other enclosed vessels^20,21^.

A way to circumvent these drawbacks is to utilize a single-objective LS, which uses the same objective for both LS formation and emission collection. In doing so, the optical complexity of the setup is reduced and the sample mounting is simplified to a compact system that can easily be incorporated on a conventional microscope stage. Crucially, the single-objective design is also ideally suited for high numerical aperture (NA) applications, including single-molecule localization microscopy (SMLM). SMLM enables super-resolution imaging of an underlying structure by separating and localizing individual fluorophores across multiple frames, overcoming the diffraction limit of light^22–26^ to uncover biological phenomena at the nanoscale^27–36^.

Certain single-objective LS approaches, such as highly laminated optical sheet (HILO)^37^ or oblique plane microscopy (OPM)^38–40^, where the sample is sectioned by tilting the illumination beam as it exits the objective, address some of the constraints imposed by two-objective designs. However, they suffer from limitations such as a limited beam thinness and coupling between effective beam thickness and LS position in the sample in the case of HILO, or a necessity for two more objectives in the detection path due to the illumination beam not being aligned with the conventional image plane in the case of OPM, limiting the photon collection efficiency. In contrast, single-objective LS illumination where the LS is reflected into the sample circumvents these drawbacks as it enables the use of a single high NA objective for high photon collection efficiency, the generation of thin LSs where the LS thickness is independent of the LS position in the sample, and provides illumination that is aligned with the sample plane being imaged. However, such reflected single-objective LS illumination designs present their own challenges, particularly the need for specialized chambers with a mechanism to reflect the LS into the sample so that the LS is aligned with the image plane. Existing chambers for single-objective LS illumination are often limited by their lack of design flexibility^41–45^, the requirement to elevate samples above the coverslip^41,45^, reliance on harsh chemicals^42^ or specialized polymers^41^, or integration into microfluidic systems prone to issues such as clogs and air bubbles^46^.

In this work, we scale and adapt our previously published versatile fabrication method for reflective micromirrors^46,47^, originally developed for small microfluidic channels, to conventional large-scale imaging chambers. Unlike our earlier approach, which embedded metallized 3D nanoprinted inserts as reflective optics within polydimethylsiloxane (PDMS) microfluidic chips prior to bonding, we now combine photolithography with macroscopic 3D printing to create molds for PDMS inserts of customizable dimensions. The metallized 3D nanoprinted micromirrors are then incorporated into these PDMS inserts, which can be easily integrated into any commercially available or user-defined imaging chamber without interfering with its functionality or drastically altering its geometry (Figure 1a). With this approach, we bypass the need for working with microfluidic channels, which are not needed for many applications and can be challenging to work with for delicate or larger biological samples, and extend our tunable fabrication scheme to a broad array of imaging chambers and applications.

**Figure 1.**
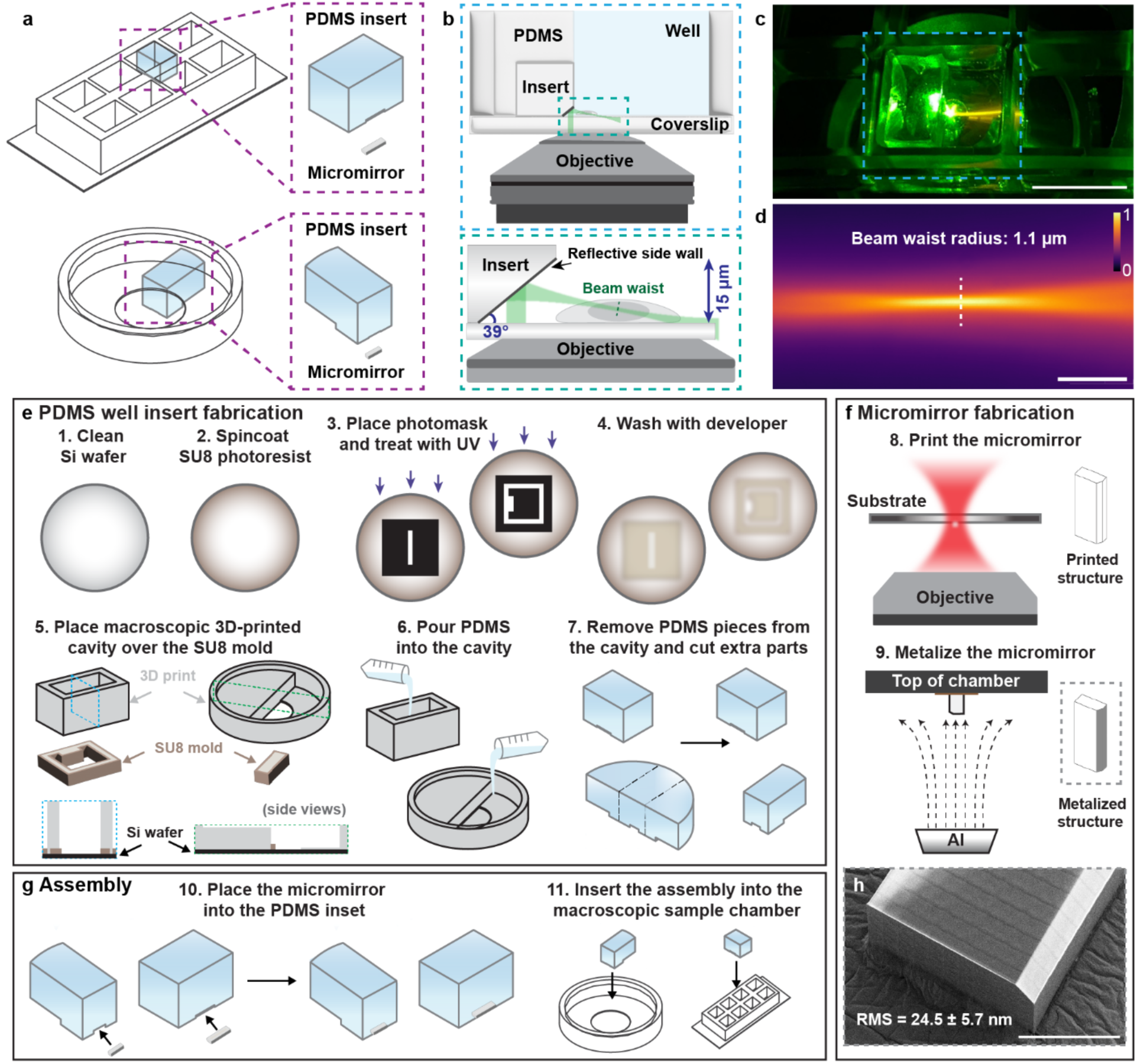
Fabrication pipeline of single-objective light sheet (LS) inserts. (a) Schematic showing the easy placement of the insert into user-defined commercially available imaging chambers. (b) Schematic of the micromirror insert system where a metalized nanoprinted insert is embedded in a PDMS piece for LS reflection into the sample. (c) Photo of the entire system on a microscope stage reflecting a single-objective LS. Scale bar is 10 mm. (d) The LS imaged in a fluorescent solution. Scale bar is 10 μm. The colorbar shows normalized intensity. (e) Schematic of the PDMS well insert fabrication pipeline. An SU8 photoresist mold is created and aligned to a 3D-printed cavity using grooves, and then PDMS is poured and cut to accommodate placement into imaging chambers. (f) Schematic of micromirror fabrication where micromirrors are printed with desired geometry using two-photon polymerization (2PP) and then metalized to be reflective using e-beam vapor deposition. (g) Schematic of insert assembly where micromirrors are placed in insets in the PDMS pieces enabled by the SU8 mold and bonded to the macroscopic sample chamber. (h) Scanning electron micrograph (SEM) of the micromirror, revealing a smooth surface after metallization. Profiled RMS is measured to be 24.5 ± 5.7 nm (mean ± standard deviation, n = 3). Scale bar is 300 μm.

Once assembled, the LS emerging from the objective is redirected by the micromirror, forming a LS at the image plane (Figure 1b,c). To ensure optimal sectioning across the imaging area, we utilize a tunable lens in the excitation path, which allows the position of the LS beam waist along the optical axis to be dynamically adjusted (Figure S1). This enables the thinnest portion of the light sheet to be precisely positioned relative to the micromirror, which is typically up to ∼50 μm from the mirror face, to coincide with the specific region of interest (Figure S2). Because the beam waist can be translated, the system maintains sectioning without requiring the cells to be cultured in a restricted or confined area near the insert. This offset ensures that the physical presence of the micromirror does not interfere with standard cell culture conditions. The LS can be tilted to enable imaging all the way down to the coverslip, circumventing the limitation imposed by the convergence angle of a Gaussian profile LS beam^15,46^. The LS tilt is directly controlled by the angle of the reflective micromirror, a parameter easily tuned through our versatile fabrication process, allowing the LS angle to be customized to suit user-specified needs. In this work, we demonstrate the inserts using a relatively thin, 1.1 μm LS (1/e^2^ beam waist radius) enabled by the high-NA single-objective design, to perform optical sectioning of mammalian cells (Figure 1d). However, given the flexibility of the fabrication method, this pipeline could easily be scaled to accommodate larger LSs for sectioning of larger systems.

The single-objective LS imaging inserts consists of two parts that are fabricated with two different processes: 1) a well insert is made with PDMS cast into a 3D-printed cavity (see Supplementary CAD files 1 and 2) placed over an SU8 mold (Figure 1e), and 2) a micromirror that is 3D nanoprinted (see Supplementary CAD files 3, 4, and 5) via two-photon polymerization^48–52^ (2PP) and metalized via e-beam vapor deposition (Figure 1f).

Well inserts are fabricated using a two-part molding approach that combines planar SU8 molds and 3D printed cavities (see Figure S3). SU8 molds are fabricated from custom-designed film photomasks^53–58^ (see Supplementary CAD files 6 and 7). Each photomask defines the lateral (XY) footprint of its corresponding imaging chamber and includes an additional region for micromirror placement. To complete the three-dimensional structure of the insert, a 3D-printed cavity is positioned on top of the SU8 mold. This cavity defines the axial morphology of the insert and features grooves that “click” onto the SU8 mold’s raised portions, enabling easy alignment of the two parts. PDMS is then poured into this assembled mold to produce a PDMS insert with consistent and reproduceable dimensions that fits precisely within the imaging chamber while maintaining an indentation for micromirror embedding.

Micromirrors are fabricated following an adapted version of the procedure described in Saliba, Gagliano, & Gustavsson^46^ and Cheng *et al*^59^. Micromirrors are first custom-designed with CAD to include desired dimensions with special attention to the angle and height of the portion used for single-objective LS reflection. These structures are digitally prepared for 3D-printing via 2PP where slicing, hatching, and printing parameters are tuned and optimized (see Table S1 for specifications). The printing material was chosen to be methacrylate-based, commercially available photoresin IP-Visio for its biocompatibility and low autofluorescence. The dimensions of the micromirror are tailored to match those of the SU8 molds that are custom-designed for the desired imaging well. The length and width of the micromirror are determined by the photomask dimensions and the height is determined based on profiled height of the SU8 photoresist layer. This precise matching ensures a seamless fit of the micromirror within the PDMS well insert. After printing, the micromirror structure is mounted on its side to prepare for metallization, ensuring only the edge containing the reflective mirror surface is coated. This prevents unnecessary reflectivity from other surfaces. The structure is then coated with 200 nm of silica, 350 nm of aluminum, and a final 5 nm layer of silica which are all deposited sequentially via e-beam vapor deposition. The initial silica layer insulates the plastic insert, protecting it from potential laser-induced heat damage, the aluminum provides reflectivity for single-objective LS reflection, and the final silica layer acts as a protective dielectric coating.

After the two parts are fabricated, they are assembled by positioning the micromirror in the correct orientation inside the indentation in the PDMS insert (Figure 1g). This “lock-and-key” fit is facilitated by a precise grove, which is defined by the SU-8 mold, acting as a physical alignment guide for the printed micromirror. To assemble the PDMS micromirror insert for single-objective LS imaging, the micromirror is placed into the indentation in the PDMS insert. By constraining the mirror within this indentation, the desired reflection geometry is consistently achieved and maintained during the assembly process, minimizing manual positional errors. Upon micromirror placement inside the PDMS groove, the assembled insert is then secured in the imaging chamber through plasma bonding, where both the glass-bottom imaging chamber and the insert are treated with air plasma. The insert is then aligned and pressed against the bottom of the chamber to form covalent bonds. These fabrication steps result in a robust and precisely aligned system, enabling effective LS imaging and cell culture within the imaging chamber. Confirmation of the function of the completed, metalized insert as a micromirror was performed with scanning electron microscopy (SEM) imaging and profilometry, revealing a smooth reflective surface with a root mean square (RMS) roughness of 24.5 ± 5.7 nm (mean ± standard deviation, n = 3) which is suitable for light reflection in microscopy applications (Figures 1h and S3). Furthermore, the assembled insert’s alignment was experimentally validated using optical microscopy, confirming that the structural grooves reliably generate the intended mirror angle (see Figure S4).

Our fabrication scheme is highly versatile, enabling generation of custom inserts for a wide range of sample chambers. Here, we demonstrate the applicability of our fabrication scheme using commercially available 8-well glass bottom chambers (Ibidi Inc.) and round glass bottom dishes (Mattek Corporation) (Figures 1 and S5). For the main results presented here, we will focus on the Ibidi chamber, but comparable performance is achieved in Mattek chambers (Figure S6). Our inserts do not interfere with the capabilities or function of the original chambers. Thus, because Ibidi chambers are non-toxic, biocompatible, and compatible with both fluorescence and transmission microscopy, we are able to perform continuous cell culture and imaging for a variety of applications. Our inserts enable easy switching between epi-illumination or transmission microscopy to HILO or single-objective LS illumination, allowing multi-modal illumination studies (Figure S1). To quantify the sectioning performance of these modalities, we first imaged 15 μm diameter fluorescent beads featuring surface-only labeling; this geometry provides a clear visual and quantitative measure of axial sectioning. While HILO illumination improves upon standard epi-illumination by tilting the excitation beam, its sectioning capability remains limited by the excitation beam’s thickness and divergence, and the coupling between sectioning plane and sectioning performance. In contrast, our micromirror-based single-objective LS maintains a thin, precisely oriented excitation plane. Line scan profiles across beads revealed that the use of LS provides an over 2x improvement in signal-to-background (SBR) compared to HILO and an over 3x improvement compared to epi-illumination (Figure 2a). This performance advantage is even more pronounced when imaging at different z-positions throughout the bead’s axial framework (Figure S7). This improved optical sectioning will also reduce photobleaching and photodamage in regions outside of the illumination volume.

**Figure 2.**
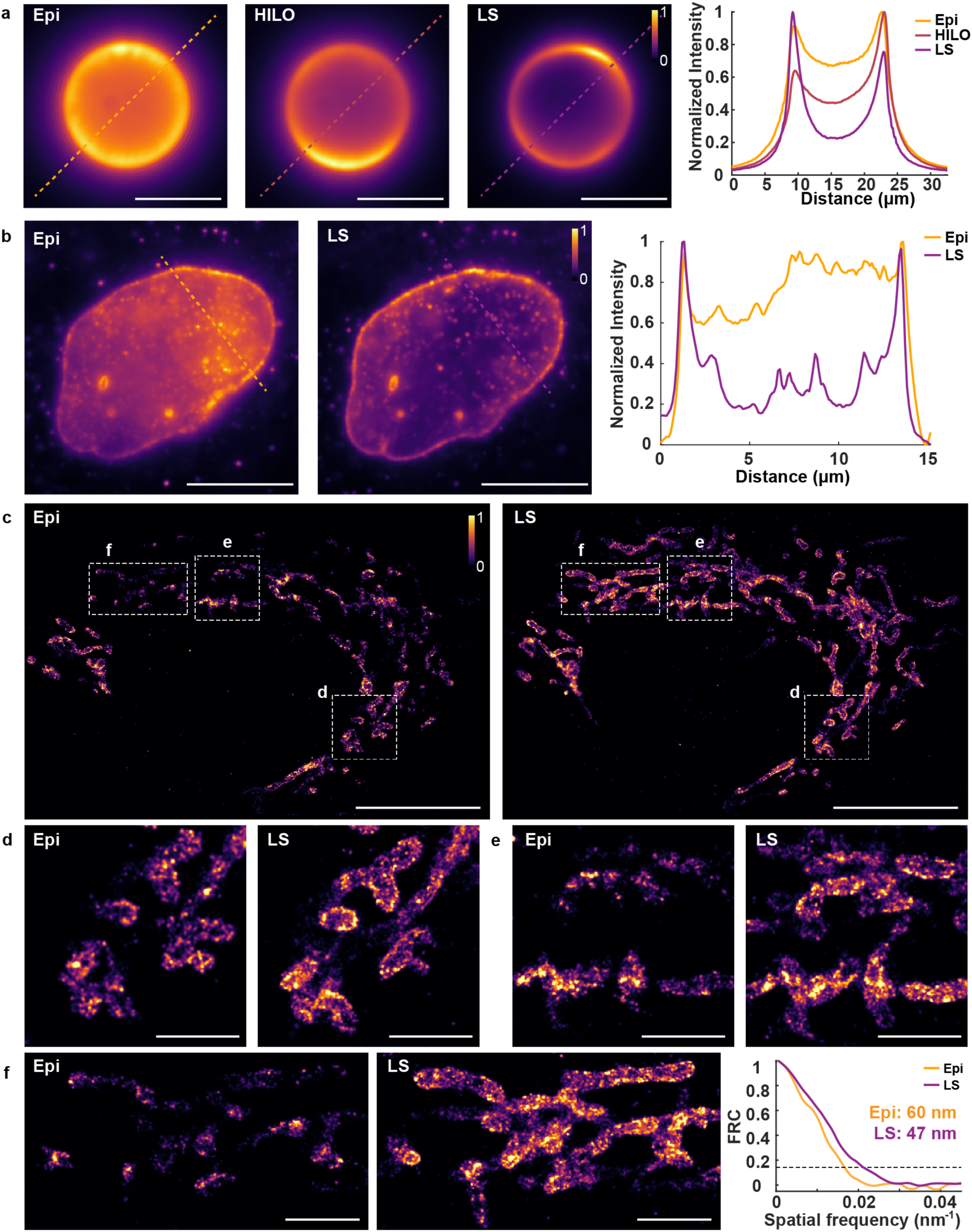
Insert-enabled single-objective LS illumination improves the signal-to-background ratio. (a) Images of a 15 μm fluorescent bead excited with epi-, highly inclined and laminated optical sheet (HILO), and LS illumination with graphs of line scans demonstrating a substantial signal-to-background (SBR) improvement with LS over both epi- and HILO modalities. Scale bars are 10 μm. (b) Diffraction-limited images of lamin B1 in a U2OS cell excited with epi- and LS illumination with graphs of line scans revealing a 4X SBR improvement with LS illumination. Scale bars are 10 μm. The colorbar shows intensity normalized independently for each image. (c-e) Single-molecule localization microscopy (SMLM) reconstructions of TOMM20 (mitochondria) using both epi- and single-objective LS illumination, revealing drastic improvements in resolvable features with LS illumination. Scale bars in (c) are 10 μm and scale bars in (d) and (e) are 1 μm. (f) Zoom in of a mitochondrial region and corresponding Fourier ring correlation (FRC) analysis curves, demonstrating an over 10 nm improvement in resolution using LS compared with epi-illumination. Scale bars are 2 μm. The colorbar shows intensity normalized to the same contrast for direct comparison between reconstructions.

To further validate the system in a biological context, we then imaged the nuclear lamina protein lamin B1 in U2OS cells with both epi- and LS illumination using our single-objective LS inserts. Line scan profiles across these images revealed an over 4X improvement in the SBR (Figures 2b and S8). This demonstrates the benefit of single-objective LS illumination for imaging cellular structures, such as the nuclear lamina, that are subject to high fluorescence background when using epi-illumination.

We then demonstrate the system for SMLM imaging using DNA points accumulation for imaging in nanoscale topography (DNA-PAINT), which involves labeling a target of interest with a short DNA oligonucleotide strand, called the docking strand, and introducing complementary dye-conjugated oligonucleotide strands, called imager strands^36,60–66^. The transient binding between the imager and docking strands achieves the temporal separation of individual fluorophores needed for SMLM. Here, we show super-resolution imaging of the mitochondrial protein TOMM20 throughout a U2OS cell using either epi- or LS illumination, where LS illumination drastically improves the resolvable features throughout the mitochondrial network (Figures 2c,d,e). LS illumination also dramatically increased the total number of detectable localizations, where we obtained a total of 714,818 detectable localizations with LS illumination compared with 282,585 localizations with epi-illumination. Fourier ring correlation (FRC) analysis^67^ was used to determine the resulting resolution of the mitochondrial structures imaged with each modality, revealing a resolution of 60 nm with epi-illumination and 47 nm with LS illumination (Figure 2f). This quantitatively confirms the improvement enabled by our inserts and LS illumination, showing an over 10 nm resolution enhancement when compared to epi-illumination (Figure 2f).

Next, multitarget single-molecule data was acquired using single-objective LS illumination enabled by reflection with the fabricated inserts. We used Exchange-PAINT, a sequential version of DNA-PAINT that utilizes orthogonal docking-imager sequences for each target to prevent crosstalk, thus enabling multitarget imaging with the same fluorophore through sequential washes^60,61^ (Table S2). We imaged the mitochondrial protein TOMM20 and the cytoskeletal protein vimentin throughout a U2OS cell, clearly resolving details of the mitochondrial network and individual filaments of vimentin (Figure 3a,b,c). The ability to image both targets was facilitated by manual sequential washes, which were easily performed because the insert does not disrupt the open-top functionality of the chamber, allowing direct access to the sample. The resulting FRC resolutions were found to be 35 nm for TOMM20 and 46 nm for vimentin, demonstrating robust performance for SMLM applications (Figures 3d and S9).

**Figure 3.**
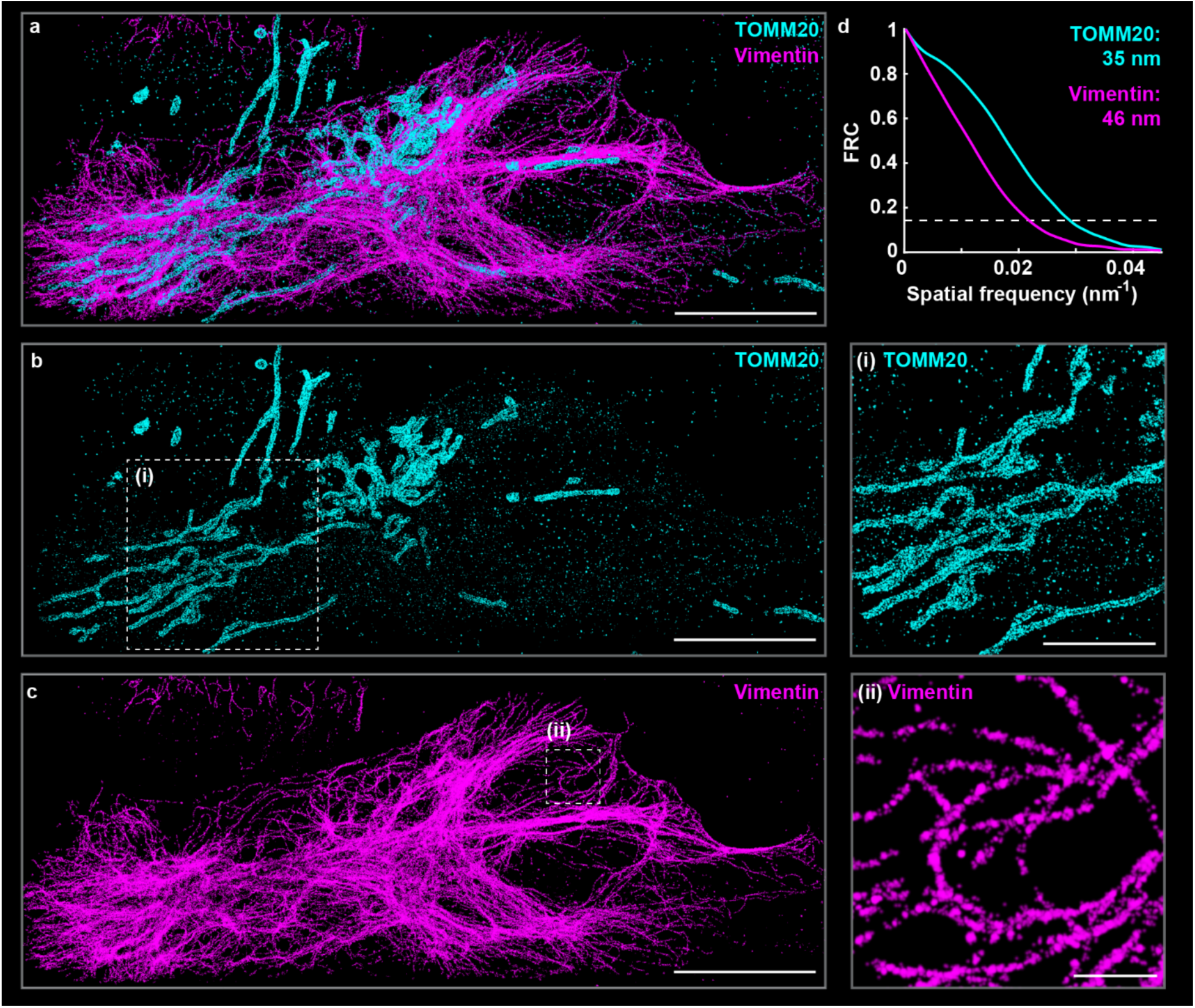
Two-target single-molecule super-resolution imaging using single-objective LS microscopy. (a) Two-target single-molecule localization microscopy reconstruction of TOMM20 (mitochondria) and vimentin in a U2OS cell and corresponding Fourier ring correlation (FRC) analysis curves. Scale bar is 10 μm. Separately rendered (b) mitochondria and (c) vimentin, with inset panels revealing details of the mitochondrial and vimentin networks. Scale bars are 10 μm for whole-cell panels, 5 μm for the inset panel in (b), and 1 μm for the inset panel in (c).

Finally, we demonstrate live-cell compatibility and the performance improvement enabled by our inserts and single-objective LS on sensitive live samples. The biocompatibility of the system allowed U2OS cells, labeled with the mitochondria-targeting live-cell dye MitoTracker, to be cultured and imaged continuously. Importantly, the entire chamber and our inserts are compatible with standard cell culture and stage-top incubators, facilitating live cell culture and imaging. Using single-objective LS illumination, we clearly resolved the dynamic movement and remodeling of the mitochondrial network (Figures 4a,b and S10). Line scan analysis revealed an improvement in the SBR of over 4X when using LS compared to epi-illumination also in the live-cell regime (Figure 4c). This drastic reduction in background is the result of optical sectioning, selective illumination of just a thin slice of the cell, when using LS illumination, which also substantially reduces phototoxicity and photobleaching compared to epi-illumination. This demonstrates the broad improvement that the implementation of our inserts can yield for various live-cell studies by enabling LS illumination in regular sample chambers.

**Figure 4.**
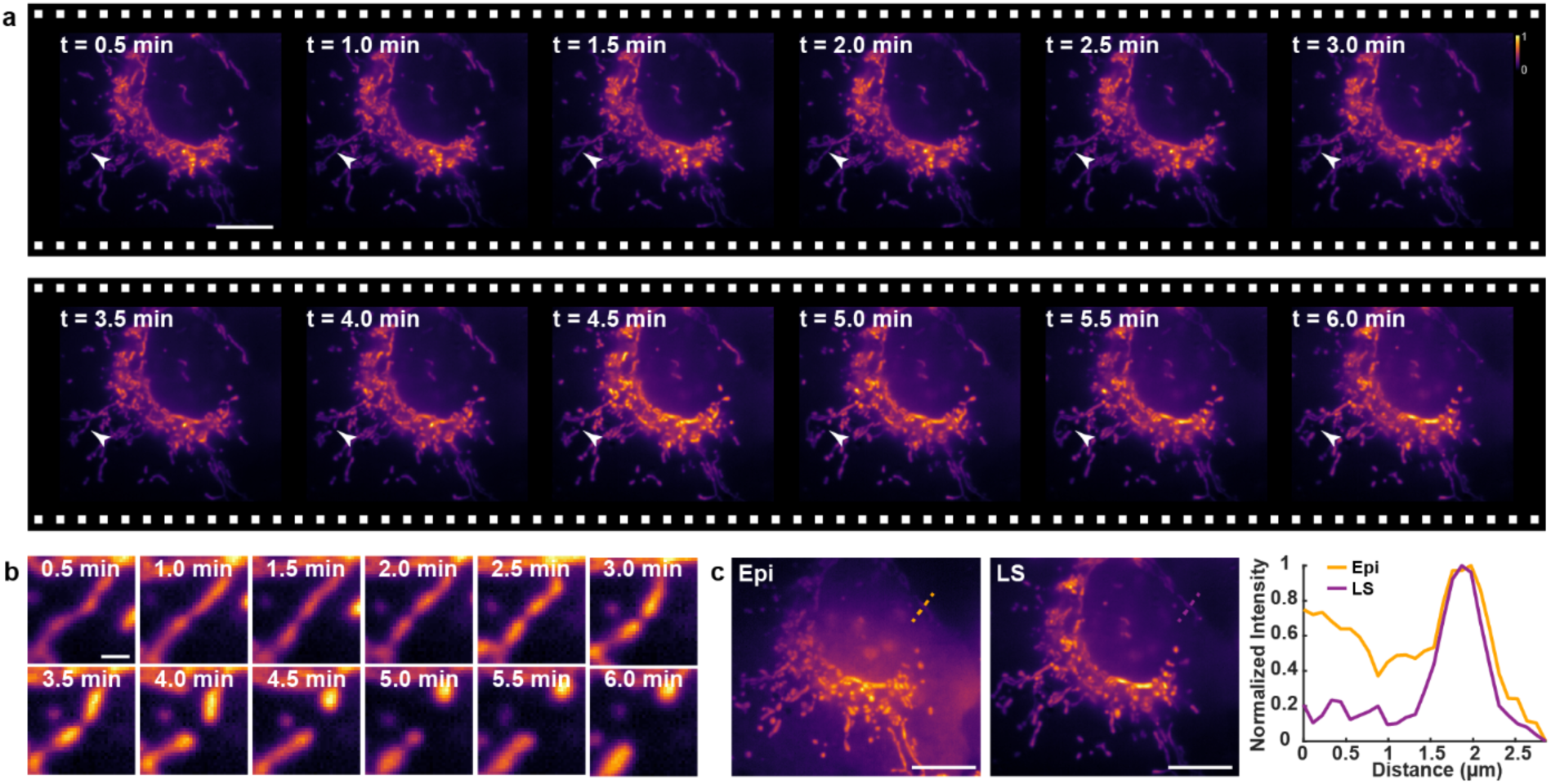
Time-lapse imaging of the mitochondrial network in live cells with single-objective LS illumination. (a) Mitochondrial network dynamics in a live U2OS cell acquired with single-objective LS time-lapse imaging. Scale bar is 10 μm. The colorbar shows intensity normalized independently for each image. (b) Zoom in revealing details of mitochondrial network remodeling over time. Scale bar is 1 μm. (c) Images of the mitochondrial network in a live cell excited with epi- and LS illumination accompanied by a graph of line scans demonstrating a 4X improvement in the signal-to-background ratio (SBR) with LS illumination. Scale bars are 10 μm.

In summary, the tunable microfabrication pipeline presented here provides a simple way to adapt conventional imaging chambers to accommodate single-objective LS illumination, which offers several benefits over conventional epi-illumination, especially for sensitive biological samples.

Because these inserts do not disrupt the original function of the imaging chambers, they can be readily used in conjunction with other illumination modalities, such epi-illumination, transmission microscopy, or total internal reflection fluorescence (TIRF) illumination^16,68^. Furthermore, they can be used with point spread function (PSF) engineering for added axial information of each emitter, enabling 3D SMLM^15,35,46,47,69–75^. Additionally, the system’s live-cell compatibility enables advanced applications such as live-cell super resolution imaging^76–79^ and single-particle tracking (SPT), particularly for investigations subject to high fluorescence background and that requires imaging away from the coverslip, such as tracking in the cell nucleus^19,80,81^, throughout the cytoplasm, or at the apical cell membrane.

While the use of 2PP reflective micromirrors has been previously demonstrated in microfluidic contexts^46,59^, our approach broadens the adaptability of the technique by decoupling implementation of the optical component from the microfluidic architecture. Unlike previous demonstrations that integrate micromirrors into microchannels, our fabrication pipeline utilizes a novel hybrid approach combining 3D 2PP nanoprinting with conventional 3D printing to create inserts compatible with conventional imaging chambers. This shift significantly increases the versatility of the technology, allowing researchers to implement single-objective LS illumination without redesigning their established cell culture or sample preparation protocols.

The insert fabrication pipeline can be easily adapted to imaging chambers of larger sizes to accommodate larger biological samples. This scaling can be simply achieved by adjusting the corresponding optics to accommodate a larger LS and by adjusting the geometry of the insert. While the fabrication time for the PDMS component remains largely independent of scale, enlarging the micromirror would increase the 3D nanoprinting duration, as print time scales with the total volume and surface area of the component being printed. However, this can be mitigated by utilizing higher speed printing modes or larger voxel sizes where high resolution is not as critical, such as for the mirror’s internal support structure, while maintaining high resolution scanning for the reflective surface. Additionally, as 3D-printing technologies advance to accommodate materials like PDMS or glass, a more streamlined fabrication pipeline may be facilitated in the future^82,83^.

Furthermore, while the fabrication process utilizes specialized equipment, including 2PP and e-beam evaporation, these tools are increasingly available through regional nanofabrication hubs. Furthermore, PDMS fabrication can be performed on a standard lab bench. To support widespread use and distribution, we have provided the 3D design files and protocols in the Supporting Information to facilitate easy replication via internal or external fabrication services.

Taken together, our insert fabrication pipeline offers an easily implemented approach for improved imaging throughout biological samples using single-objective LS illumination in conventional imaging chambers.

## ASSOCIATED CONTENT

### Supporting Information

The Supporting Information is available free of charge.

- The Supporting Information pdf file contains further details on insert design and fabrication, sample preparation, image acquisition, and data analysis and rendering, as well as figures and tables with supporting data and further statistics from imaging acquisitions.
- Supplementary CAD files for 3D printing of PDMS insert cavities (SI_CAD_1-2), for nanoprinting of the micromirror inserts (SI_CAD_3-5), and of the photomasks for SU8 mold fabrication (SI_CAD_6-7) can be found on GitHub (https://github.com/Gustavsson-Lab/LS-fab).

## AUTHOR INFORMATION

### Author Contributions

N.S. conceived the project. N.S, S.C., and A.-K.G. directed the project. N.S. and S.C. optimized the fabrication pipeline, prepared samples, and analyzed and rendered data. P.J. constructed the imaging platform. N.S., S.C., and P.J. acquired data. N.S., S.C., and A.-K.G wrote the manuscript and prepared figures with input from all other authors.

## Notes

N.S., S.C., and A.-K.G. are listed as inventors on patent applications filed by Rice University that describe the fabrication process detailed in this manuscript.

## ACKNOWLEDGMENT

The authors thank Yuya Nakatani, Gabriella Gagliano, and Ivana Hsyung for assistance with sample labeling, optical alignment, and preliminary prototypes. Insert fabrication in this work were fabricated, imaged, and profiled using resources and equipment available through the Shared Equipment Authority at Rice University.

## REFERENCES

(1) Power, R. M.; Huisken, J. A Guide to Light-Sheet Fluorescence Microscopy for Multiscale Imaging. Nat. Methods 2017, 14 (4), 360–373. 10.1038/nmeth.4224.

(2) Gustavsson, A.-K.; Petrov, P. N.; Moerner, W. E. Light Sheet Approaches for Improved Precision in 3D Localization-Based Super-Resolution Imaging in Mammalian Cells [Invited]. Opt. Express 2018, 26 (10), 13122. 10.1364/OE.26.013122.

(3) Wan, Y.; McDole, K.; Keller, P. J. Light-Sheet Microscopy and Its Potential for Understanding Developmental Processes. Annu. Rev. Cell Dev. Biol. 2019, 35 (1), 655–681. 10.1146/annurev-cellbio-100818-125311.

(4) Gagliano, G.; Nelson, T.; Saliba, N.; Vargas-Hernández, S.; Gustavsson, A.-K. Light Sheet Illumination for 3D Single-Molecule Super-Resolution Imaging of Neuronal Synapses. Front. Synaptic Neurosci. 2021, 13, 761530. 10.3389/fnsyn.2021.761530.

(5) Cheng, S.; Nakatani, Y.; Gagliano, G.; Saliba, N.; Gustavsson, A.-K. Light Sheet Illumination in Single-Molecule Localization Microscopy for Imaging of Cellular Architectures and Molecular Dynamics. npj Imaging 2024, 2 (1), 49. 10.1038/s44303-024-00057-9.

(6) Kramer, S. N.; Antarasen, J.; Reinholt, C. R.; Kisley, L. A Practical Guide to Light-Sheet Microscopy for Nanoscale Imaging: Looking beyond the Cell. Journal of Applied Physics 2024, 136 (9), 091101. 10.1063/5.0218262.

(7) Voie, A. H.; Burns, D. H.; Spelman, F. A. Orthogonal-Plane Fluorescence Optical Sectioning: Three-Dimensional Imaging of Macroscopic Biological Specimens. J. Microsc. 1993, 170 (3), 229–236. 10.1111/j.1365-2818.1993.tb03346.x.

(8) Huisken, J.; Swoger, J.; Del Bene, F.; Wittbrodt, J.; Stelzer, E. H. K. Optical Sectioning Deep inside Live Embryos by Selective Plane Illumination Microscopy. Science 2004, 305 (5686), 1007. 10.1126/science.1100035.

(9) Cella Zanacchi, F.; Lavagnino, Z.; Perrone Donnorso, M.; Del Bue, A.; Furia, L.; Faretta, M.; Diaspro, A. Live-Cell 3D Super-Resolution Imaging in Thick Biological Samples. Nat. Methods 2011, 8 (12), 1047–1049. 10.1038/nmeth.1744.

(10) Gebhardt, J. C. M.; Suter, D. M.; Roy, R.; Zhao, Z. W.; Chapman, A. R.; Basu, S.; Maniatis, T.; Xie, X. S. Single-Molecule Imaging of Transcription Factor Binding to DNA in Live Mammalian Cells. Nat. Methods 2013, 10 (5), 421–426. 10.1038/nmeth.2411.

(11) Hu, Y. S.; Zhu, Q.; Elkins, K.; Tse, K.; Li, Y.; Fitzpatrick, J. A. J.; Verma, I. M.; Cang, H. Light-Sheet Bayesian Microscopy Enables Deep-Cell Super-Resolution Imaging of Heterochromatin in Live Human Embryonic Stem Cells. Opt. Nanoscopy 2013, 2 (1), 7. 10.1186/2192-2853-2-7.

(12) Zhao, Z. W.; Roy, R.; Gebhardt, J. C. M.; Suter, D. M.; Chapman, A. R.; Xie, X. S. Spatial Organization of RNA Polymerase II inside a Mammalian Cell Nucleus Revealed by Reflected Light-Sheet Superresolution Microscopy. Proceedings of the National Academy of Sciences 2014, 111 (2), 681–686. 10.1073/pnas.1318496111.

(13) Greiss, F.; Deligiannaki, M.; Jung, C.; Gaul, U.; Braun, D. Single-Molecule Imaging in Living Drosophila Embryos with Reflected Light-Sheet Microscopy. Biophys. J. 2016, 110 (4), 939–946. 10.1016/j.bpj.2015.12.035.

(14) Fadero, T. C.; Gerbich, T. M.; Rana, K.; Suzuki, A.; DiSalvo, M.; Schaefer, K. N.; Heppert, J. K.; Boothby, T. C.; Goldstein, B.; Peifer, M.; Allbritton, N. L.; Gladfelter, A. S.; Maddox, A. S.; Maddox, P. S. LITE Microscopy: Tilted Light-Sheet Excitation of Model Organisms Offers High Resolution and Low Photobleaching. J. Cell Biol. 2018, 217 (5), 1869–1882. 10.1083/jcb.201710087.

(15) Gustavsson, A.-K.; Petrov, P. N.; Lee, M. Y.; Shechtman, Y.; Moerner, W. E. 3D Single-Molecule Super-Resolution Microscopy with a Tilted Light Sheet. Nat Commun 2018, 9 (1), 123. 10.1038/s41467-017-02563-4.

(16) Nelson, T.; Vargas-Hernández, S.; Freire, M.; Cheng, S.; Gustavsson, A.-K. Multimodal Illumination Platform for 3D Single-Molecule Super-Resolution Imaging throughout Mammalian Cells. Biomed. Opt. Express 2024, 15 (5), 3050–3063. 10.1364/BOE.521362.

(17) Chen, B.-C.; Legant, W. R.; Wang, K.; Shao, L.; Milkie, D. E.; Davidson, M. W.; Janetopoulos, C.; Wu, X. S.; Hammer, J. A.; Liu, Z.; English, B. P.; Mimori-Kiyosue, Y.; Romero, D. P.; Ritter, A. T.; Lippincott-Schwartz, J.; Fritz-Laylin, L.; Mullins, R. D.; Mitchell, D. M.; Bembenek, J. N.; Reymann, A.-C.; Böhme, R.; Grill, S. W.; Wang, J. T.; Seydoux, G.; Tulu, U. S.; Kiehart, D. P.; Betzig, E. Lattice Light-Sheet Microscopy: Imaging Molecules to Embryos at High Spatiotemporal Resolution. Science 2014, 346 (6208), 1257998. 10.1126/science.1257998.

(18) Legant, W. R.; Shao, L.; Grimm, J. B.; Brown, T. A.; Milkie, D. E.; Avants, B. B.; Lavis, L. D.; Betzig, E. High-Density Three-Dimensional Localization Microscopy across Large Volumes. Nat Methods 2016, 13 (4), 359–365. 10.1038/nmeth.3797.

(19) Gustavsson, A.-K.; Ghosh, R. P.; Petrov, P. N.; Liphardt, J. T.; Moerner, W. E. Fast and Parallel Nanoscale Three-Dimensional Tracking of Heterogeneous Mammalian Chromatin Dynamics. Mol. Biol. Cell 2022, 33 (6), 1–11. 10.1091/mbc.E21-10-0514.

(20) Zhang, J.; Zhang, M.; Wang, Y.; Donarski, E.; Gahlmann, A. Optically Accessible Microfluidic Flow Channels for Noninvasive High-Resolution Biofilm Imaging Using Lattice Light Sheet Microscopy. J. Phys. Chem. B 2021, 125 (44), 12187–12196. 10.1021/acs.jpcb.1c07759.

(21) Moore, R. P.; O’Shaughnessy, E. C.; Shi, Y.; Nogueira, A. T.; Heath, K. M.; Hahn, K. M.; Legant, W. R. A Multi-Functional Microfluidic Device Compatible with Widefield and Light Sheet Microscopy. Lab Chip 2021, 22 (1), 136–147. 10.1039/D1LC00600B.

(22) Sharonov, A.; Hochstrasser, R. M. Wide-Field Subdiffraction Imaging by Accumulated Binding of Diffusing Probes. Proceedings of the National Academy of Sciences 2006, 103 (50), 18911–18916. 10.1073/pnas.0609643104.

(23) Betzig, E.; Patterson, G. H.; Sougrat, R.; Lindwasser, O. W.; Olenych, S.; Bonifacino, J. S.; Davidson, M. W.; Lippincott-Schwartz, J.; Hess, H. F. Imaging Intracellular Fluorescent Proteins at Nanometer Resolution. Science 2006, 313 (5793), 1642–1645. 10.1126/science.1127344.

(24) Rust, M. J.; Bates, M.; Zhuang, X. Sub-Diffraction-Limit Imaging by Stochastic Optical Reconstruction Microscopy (STORM). Nat Methods 2006, 3 (10), 793–796. 10.1038/nmeth929.

(25) Hess, S. T.; Girirajan, T. P. K.; Mason, M. D. Ultra-High Resolution Imaging by Fluorescence Photoactivation Localization Microscopy. Biophys. J. 2006, 91 (11), 4258–4272. 10.1529/biophysj.106.091116.

(26) Weiss, L. E.; Love, J. F.; Yoon, J.; Comerci, C. J.; Milenkovic, L.; Kanie, T.; Jackson, P. K.; Stearns, T.; Gustavsson, A.-K. Chapter 4 - Single-Molecule Imaging in the Primary Cilium. In Methods Cell Biol; Bravo-San Pedro, J. M., Galluzzi, L., Eds.; Academic Press, 2023; Vol. 176, pp 59–83. 10.1016/bs.mcb.2023.01.003.

(27) Xu, K.; Babcock, H. P.; Zhuang, X. Dual-Objective STORM Reveals Three-Dimensional Filament Organization in the Actin Cytoskeleton. Nat Methods 2012, 9 (2), 185–188. 10.1038/nmeth.1841.

(28) Andronov, L.; Han, M.; Zhu, Y.; Balaji, A.; Roy, A. R.; Barentine, A. E. S.; Patel, P.; Garhyan, J.; Qi, L. S.; Moerner, W. E. Nanoscale Cellular Organization of Viral RNA and Proteins in SARS-CoV-2 Replication Organelles. Nat Commun 2024, 15 (1), 4644. 10.1038/s41467-024-48991-x.

(29) Chowdhury, P.; Wang, X.; Han, R. I.; Motrapu, M.; Boice, A. G.; Nakatani, Y.; Vargas-Hernandez, S.; Love, J. F.; Chew, C.; Grimm, S. L.; Mezquita, D.; Mason, F. M.; Martinez, E. D.; Coarfa, C.; Walker, C. L.; Gustavsson, A.; Dere, R. Lysine Demethylase 4A Is a Centrosome-associated Protein Required for Centrosome Integrity and Genomic Stability. The FEBS Journal 2026, 293 (2), 396–417. 10.1111/febs.70240.

(30) Van Veen, S.; Kuipers, M.; Nolte-’t Hoen, E.; Albertazzi, L. Uncovering Subpopulations in Extracellular Vesicles via Multiplexed Spectral PAINT Super-Resolution Microscopy. Nano Lett. 2026, 26 (4), 1163–1170. 10.1021/acs.nanolett.5c03118.

(31) Winkler, M.; Mack, T. G. A.; Eickholt, B. J.; Schmoranzer, J.; Burn, G. L.; Gimber, N. Nanoscale Mapping Reveals Periodic Organization of Neutrophil Extracellular Trap Proteins. Nano Lett. 2026, acs.nanolett.5c05175. 10.1021/acs.nanolett.5c05175.

(32) Appelhans, T.; Richter, C. P.; Wilkens, V.; Hess, S. T.; Piehler, J.; Busch, K. B. Nanoscale Organization of Mitochondrial Microcompartments Revealed by Combining Tracking and Localization Microscopy. Nano Lett. 2012, 12 (2), 610–616. 10.1021/nl203343a.

(33) Velas, L.; Brameshuber, M.; Huppa, J. B.; Kurz, E.; Dustin, M. L.; Zelger, P.; Jesacher, A.; Schütz, G. J. Three-Dimensional Single Molecule Localization Microscopy Reveals the Topography of the Immunological Synapse at Isotropic Precision below 15 Nm. Nano Lett. 2021, 21 (21), 9247–9255. 10.1021/acs.nanolett.1c03160.

(34) Kaplan, C.; Jing, B.; Winterflood, C. M.; Bridges, A. A.; Occhipinti, P.; Schmied, J.; Grinhagens, S.; Gronemeyer, T.; Tinnefeld, P.; Gladfelter, A. S.; Ries, J.; Ewers, H. Absolute Arrangement of Subunits in Cytoskeletal Septin Filaments in Cells Measured by Fluorescence Microscopy. Nano Lett. 2015, 15 (6), 3859–3864. 10.1021/acs.nanolett.5b00693.

(35) Möckl, L.; Pedram, K.; Roy, A. R.; Krishnan, V.; Gustavsson, A.-K.; Dorigo, O.; Bertozzi, C. R.; Moerner, W. E. Quantitative Super-Resolution Microscopy of the Mammalian Glycocalyx. Dev. Cell 2019, 50 (1), 57–72. 10.1016/j.devcel.2019.04.035.

(36) Unterauer, E. M.; Shetab Boushehri, S.; Jevdokimenko, K.; Masullo, L. A.; Ganji, M.; Sograte-Idrissi, S.; Kowalewski, R.; Strauss, S.; Reinhardt, S. C. M.; Perovic, A.; Marr, C.; Opazo, F.; Fornasiero, E. F.; Jungmann, R. Spatial Proteomics in Neurons at Single-Protein Resolution. Cell 2024, 187 (7), 1785–1800.e16. 10.1016/j.cell.2024.02.045.

(37) Tokunaga, M.; Imamoto, N.; Sakata-Sogawa, K. Highly Inclined Thin Illumination Enables Clear Single-Molecule Imaging in Cells. Nat Methods 2008, 5 (2), 159–161. 10.1038/nmeth1171.

(38) Dunsby, C. Optically Sectioned Imaging by Oblique Plane Microscopy. Opt. Express 2008, 16 (11), 20306–20316. 10.1364/OE.16.020306.

(39) Sparks, H.; Dent, L.; Bakal, C.; Behrens, A.; Salbreux, G.; Dunsby, C.; Dunsby, C. Dual-View Oblique Plane Microscopy (dOPM). Biomed. Opt. Express, BOE 2020, 11 (12), 7204–7220. 10.1364/BOE.409781.

(40) Chen, B.; Chang, B.-J.; Zhou, F. Y.; Daetwyler, S.; Sapoznik, E.; Nanes, B. A.; Terrazas, I.; Gihana, G. M.; Castro, L. P.; Chan, I. S.; Conacci-Sorrell, M.; Dean, K. M.; Millett-Sikking, A.; York, A. G.; Fiolka, R. Increasing the Field-of-View in Oblique Plane Microscopy via Optical Tiling. Biomed. Opt. Express, BOE 2022, 13 (11), 5616–5627. 10.1364/BOE.467969.

(41) Galland, R.; Grenci, G.; Aravind, A.; Viasnoff, V.; Studer, V.; Sibarita, J.-B. 3D High- and Super-Resolution Imaging Using Single-Objective SPIM. Nat. Methods 2015, 12 (7), 641–644. 10.1038/nmeth.3402.

(42) Meddens, M. B. M.; Liu, S.; Finnegan, P. S.; Edwards, T. L.; James, C. D.; Lidke, K. A. Single Objective Light-Sheet Microscopy for High-Speed Whole-Cell 3D Super-Resolution. Biomed. Opt. Express 2016, 7 (6), 2219. 10.1364/BOE.7.002219.

(43) Ponjavic, A.; Ye, Y.; Laue, E.; Lee, S. F.; Klenerman, D. Sensitive Light-Sheet Microscopy in Multiwell Plates Using an AFM Cantilever. Biomed. Opt. Express 2018, 9 (12), 5863–5880. 10.1364/BOE.9.005863.

(44) Beghin, A.; Kechkar, A.; Butler, C.; Levet, F.; Cabillic, M.; Rossier, O.; Giannone, G.; Galland, R.; Choquet, D.; Sibarita, J.-B. Localization-Based Super-Resolution Imaging Meets High-Content Screening. Nat Methods 2017, 14 (12), 1184–1190. 10.1038/nmeth.4486.

(45) Galgani, T.; Fedala, Y.; Zapata, R.; Caccianini, L.; Viasnoff, V.; Sibarita, J.-B.; Galland, R.; Dahan, M.; Hajj, B. Selective Volumetric Excitation and Imaging for Single Molecule Localization Microscopy in Multicellular Systems. bioRxiv December 3, 2022, p 2022.12.02.518828. 10.1101/2022.12.02.518828.

(46) Saliba, N.; Gagliano, G.; Gustavsson, A.-K. Whole-Cell Multi-Target Single-Molecule Super-Resolution Imaging in 3D with Microfluidics and a Single-Objective Tilted Light Sheet. Nature Communications 2024, 15 (1), 10187. 10.1038/s41467-024-54609-z.

(47) Cheng, S.; Saliba, N.; Gagliano, G.; Joshi, P.; Gustavsson, A.-K. Single-Objective Lattice Light Sheet Microscopy with Microfluidics for Single-Molecule Super-Resolution Imaging of Mammalian Cells. bioRxiv July 23, 2025. 10.1101/2025.07.18.665606.

(48) Lee, K.; Yang, D.; Park, S. H.; Kim, R. H. Recent Developments in the Use of Two-photon Polymerization in Precise 2D and 3D Microfabrications. Polymers for Advanced Techs 2006, 17 (2), 72–82. 10.1002/pat.664.

(49) Boudene, I.; Bougdid, Y. Two-Photon Polymerization-Assisted 3D Laser Nanoprinting: From Fundamentals to Modern Applications. J. Mater. Chem. C 2025, 13 (36), 18597–18630. 10.1039/D5TC02037A.

(50) Zhang, L.; Wang, C.; Zhang, C.; Xue, Y.; Ye, Z.; Xu, L.; Hu, Y.; Li, J.; Chu, J.; Wu, D. High-Throughput Two-Photon 3D Printing Enabled by Holographic Multi-Foci High-Speed Scanning. Nano Lett. 2024, 24 (8), 2671–2679. 10.1021/acs.nanolett.4c00505.

(51) Khan, S. B.; Irfan, S.; Zhang, Z.; Yuan, W. Redefining Medical Applications with Safe and Sustainable 3D Printing. ACS Appl. Bio Mater. 2025, 8 (8), 6470–6525. 10.1021/acsabm.4c01923.

(52) Claeyssens, F.; Hasan, E. A.; Gaidukeviciute, A.; Achilleos, D. S.; Ranella, A.; Reinhardt, C.; Ovsianikov, A.; Shizhou, X.; Fotakis, C.; Vamvakaki, M.; Chichkov, B. N.; Farsari, M. Three-Dimensional Biodegradable Structures Fabricated by Two-Photon Polymerization. Langmuir 2009, 25 (5), 3219–3223. 10.1021/la803803m.

(53) Gustavsson, A.-K.; van Niekerk, D. D.; Adiels, C. B.; du Preez, F. B.; Goksör, M.; Snoep, J. L. Sustained Glycolytic Oscillations in Individual Isolated Yeast Cells. FEBS J. 2012, 279 (16), 2837–2847. 10.1111/j.1742-4658.2012.08639.x.

(54) Gustavsson, A.-K. Heterogeneity of Glycolytic Oscillatory Behaviour in Individual Yeast Cells. FEBS Lett. 2014, 588, 3–7.

(55) Gustavsson, A.; Banaeiyan, A. A.; Niekerk, D. D.; Snoep, J. L.; Adiels, C. B.; Goksör, M. Studying Glycolytic Oscillations in Individual Yeast Cells by Combining Fluorescence Microscopy with Microfluidics and Optical Tweezers. Curr. Prot. Cell Biol. 2019, 82 (1), 1–26.

(56) van Niekerk, D. D.; Gustavsson, A.-K.; Mojica-Benavides, M.; Adiels, C. B.; Goksör, M.; Snoep, J. L. Phosphofructokinase Controls the Acetaldehyde-Induced Phase Shift in Isolated Yeast Glycolytic Oscillators. Biochem. J. 2019, 476 (2), 353–363. 10.1042/BCJ20180757.

(57) Gustavsson, A.-K.; Adiels, C. B.; Mehlig, B.; Goksör, M. Entrainment of Heterogeneous Glycolytic Oscillations in Single Cells. Sci. Rep. 2015, 5 (9404), 1–7. 10.1038/srep09404.

(58) Gustavsson, A.-K.; van Niekerk, D. D.; Adiels, C. B.; Kooi, B.; Goksör, M.; Snoep, J. L. Allosteric Regulation of Phosphofructokinase Controls the Emergence of Glycolytic Oscillations in Isolated Yeast Cells. FEBS J. 2014, 281, 2784–2793.

(59) Cheng, S.; Saliba, N.; Gagliano, G.; Joshi, P.; Gustavsson, A.-K. Single-Objective Lattice Light Sheet Microscopy with Microfluidics for Single-Molecule Super-Resolution Imaging of Mammalian Cells. ACS Photonics 2026, 13 (1), 249–262. 10.1021/acsphotonics.5c02201.

(60) Jungmann, R.; Avendaño, M. S.; Woehrstein, J. B.; Dai, M.; Shih, W. M.; Yin, P. Multiplexed 3D Cellular Super-Resolution Imaging with DNA-PAINT and Exchange-PAINT. Nat Methods 2014, 11 (3), 313–318. 10.1038/nmeth.2835.

(61) Schnitzbauer, J.; Strauss, M. T.; Schlichthaerle, T.; Schueder, F.; Jungmann, R. Super-Resolution Microscopy with DNA-PAINT. Nat. Protoc. 2017, 12 (6), 1198–1228. 10.1038/nprot.2017.024.

(62) Steen, P. R.; Unterauer, E. M.; Masullo, L. A.; Kwon, J.; Perovic, A.; Jevdokimenko, K.; Opazo, F.; Fornasiero, E. F.; Jungmann, R. The DNA-PAINT Palette: A Comprehensive Performance Analysis of Fluorescent Dyes. Nat Methods 2024, 21 (9), 1755–1762. 10.1038/s41592-024-02374-8.

(63) Wade, O. K.; Woehrstein, J. B.; Nickels, P. C.; Strauss, S.; Stehr, F.; Stein, J.; Schueder, F.; Strauss, M. T.; Ganji, M.; Schnitzbauer, J.; Grabmayr, H.; Yin, P.; Schwille, P.; Jungmann, R. 124-Color Super-Resolution Imaging by Engineering DNA-PAINT Blinking Kinetics. Nano Lett. 2019, 19 (4), 2641–2646. 10.1021/acs.nanolett.9b00508.

(64) Stein, J.; Stehr, F.; Schueler, P.; Blumhardt, P.; Schueder, F.; Mücksch, J.; Jungmann, R.; Schwille, P. Toward Absolute Molecular Numbers in DNA-PAINT. Nano Lett. 2019, 19 (11), 8182–8190. 10.1021/acs.nanolett.9b03546.

(65) Auer, A.; Strauss, M. T.; Schlichthaerle, T.; Jungmann, R. Fast, Background-Free DNA-PAINT Imaging Using FRET-Based Probes. Nano Lett. 2017, 17 (10), 6428–6434. 10.1021/acs.nanolett.7b03425.

(66) Schueder, F.; Rivera-Molina, F.; Su, M.; Marin, Z.; Kidd, P.; Rothman, J. E.; Toomre, D.; Bewersdorf, J. Unraveling Cellular Complexity with Transient Adapters in Highly Multiplexed Super-Resolution Imaging. Cell 2024, 187 (7), 1769–1784.e18. 10.1016/j.cell.2024.02.033.

(67) Nieuwenhuizen, R. P. J.; Lidke, K. A.; Bates, M.; Puig, D. L.; Grünwald, D.; Stallinga, S.; Rieger, B. Measuring Image Resolution in Optical Nanoscopy. Nat Methods 2013, 10 (6), 557–562. 10.1038/nmeth.2448.

(68) Axelrod, D. Total Internal Reflection Fluorescence Microscopy in Cell Biology. Traffic 2001, 2 (11), 764–774. 10.1034/j.1600-0854.2001.21104.x.

(69) Pavani, S. R. P.; Thompson, M. A.; Biteen, J. S.; Lord, S. J.; Liu, N.; Twieg, R. J.; Piestun, R.; Moerner, W. E. Three-Dimensional, Single-Molecule Fluorescence Imaging beyond the Diffraction Limit by Using a Double-Helix Point Spread Function. Proc. Natl. Acad. Sci. USA 2009, 106 (9), 2995–2999. 10.1073/pnas.0900245106.

(70) Nakatani, Y.; Gaumer, S.; Shechtman, Y.; Gustavsson, A.-K. Long Axial-Range Double-Helix Point Spread Functions for 3D Volumetric Super-Resolution Imaging. in press, Journal of Physical Chemistry B 2024. 10.1101/2024.07.31.605907.

(71) Shechtman, Y.; Weiss, L. E.; Backer, A. S.; Sahl, S. J.; Moerner, W. E. Precise Three-Dimensional Scan-Free Multiple-Particle Tracking over Large Axial Ranges with Tetrapod Point Spread Functions. Nano Lett. 2015, 15 (6), 4194–4199. 10.1021/acs.nanolett.5b01396.

(72) Bennett, H. W.; Gustavsson, A.-K.; Bayas, C. A.; Petrov, P. N.; Mooney, N.; Moerner, W. E.; Jackson, P. K. Novel Fibrillar Structure in the Inversin Compartment of Primary Cilia Revealed by 3D Single-Molecule Superresolution Microscopy. Mol. Biol. Cell 2020, 31 (7), 619–639. 10.1091/mbc.E19-09-0499.

(73) Bayas, C. A.; Diezmann, A. von; Gustavsson, A.-K.; Moerner, W. E. Easy-DHPSF 2.0: Open-Source Software for Three-Dimensional Localization and Two-Color Registration of Single Molecules with Nanoscale Accuracy. Prot. Exch. 2019, 1–28. 10.21203/rs.2.9151/v2.

(74) Kanie, T.; Liu, B.; Love, J. F.; Fisher, S. D.; Gustavsson, A.-K.; Jackson, P. K. A Hierarchical Pathway for Assembly of the Distal Appendages That Organize Primary Cilia. eLife 2025, 14, e85999. 10.7554/eLife.85999.

(75) Chowdhury, P.; Wang, X.; Han, R. I.; Motrapu, M.; Boice, A. G.; Nakatani, Y.; Vargas-Hernandez, S.; Love, J. F.; Chew, C.; Grimm, S. L.; Mezquita, D.; Mason, F. M.; Martinez, E. D.; Coarfa, C.; Walker, C. L.; Gustavsson, A.; Dere, R. Lysine Demethylase 4A Is a Centrosome-associated Protein Required for Centrosome Integrity and Genomic Stability. The FEBS Journal 2025, febs.70240. 10.1111/febs.70240.

(76) Eklund, A. S.; Ganji, M.; Gavins, G.; Seitz, O.; Jungmann, R. Peptide-PAINT Super-Resolution Imaging Using Transient Coiled Coil Interactions. Nano Lett. 2020, 20 (9), 6732–6737. 10.1021/acs.nanolett.0c02620.

(77) Saguy, A.; Alalouf, O.; Opatovski, N.; Jang, S.; Heilemann, M.; Shechtman, Y. DBlink: Dynamic Localization Microscopy in Super Spatiotemporal Resolution via Deep Learning. Nature Methods 2023, 20 (12), 1939–1948. 10.1038/s41592-023-01966-0.

(78) Oi, C.; Gidden, Z.; Holyoake, L.; Kantelberg, O.; Mochrie, S.; Horrocks, M. H.; Regan, L. LIVE-PAINT Allows Super-Resolution Microscopy inside Living Cells Using Reversible Peptide-Protein Interactions. Commun Biol 2020, 3 (1), 458. 10.1038/s42003-020-01188-6.

(79) Bhaskar, H.; Kleinjan, D.; Oi, C.; Gidden, Z.; Rosser, S. J.; Horrocks, M. H.; Regan, L. Live-cell Super-resolution Imaging of Actin Using LIFEACT -14 with a PAINT -based Approach. Protein Science 2023, 32 (2), e4558. 10.1002/pro.4558.

(80) Raterink, A.; Ghosh, R. P.; Yang, L.; Shi, Q.; Nguyen, M. K.; Hilton, I. B.; Liphardt, J. T.; Gustavsson, A.-K. Robust Fluorescent Labeling and Tracking of Endogenous Non-Repetitive Genomic Loci. bioRxiv August 27, 2025. 10.1101/2025.08.22.671818.

(81) Li, J.; Liu, S.; Kim, S.; Goell, J.; Drum, Z. A.; Flores, J. P.; Ma, A. J.; Mahata, B.; Escobar, M.; Raterink, A.; Ahn, J. H.; Terán, E. R.; Guerra-Resendez, R. S.; Zhou, Y.; Yu, B.; Diehl, M. R.; Wang, G. G.; Gustavsson, A.-K.; Phanstiel, D. H.; Hilton, I. B. Biomolecular Condensation of Human IDRs Initiates Endogenous Transcription via Intrachromosomal Looping or High-Density Promoter Localization. Nucleic Acids Research 2025, 53 (4), gkaf056. 10.1093/nar/gkaf056.

(82) Du, K.; Basuki, J.; Glattauer, V.; Mesnard, C.; Nguyen, A. T.; Alexander, D. L. J.; Hughes, T. C. Digital Light Processing 3D Printing of PDMS-Based Soft and Elastic Materials with Tunable Mechanical Properties. ACS Appl. Polym. Mater. 2021, 3 (6), 3049–3059. 10.1021/acsapm.1c00260.

(83) Kotz, F.; Quick, A. S.; Risch, P.; Martin, T.; Hoose, T.; Thiel, M.; Helmer, D.; Rapp, B. E. Two-Photon Polymerization of Nanocomposites for the Fabrication of Transparent Fused Silica Glass Microstructures. Advanced Materials 2021, 33 (9), 2006341. 10.1002/adma.202006341.

