## Supplementary Information for "Versatile and Scalable Reflective Micromirrors for Single-Objective Light Sheet Microscopy"

This supplementary file includes:

- Supplementary Materials and Methods
    - PDMS well insert fabrication
    - Micromirror fabrication
    - SEM imaging
    - Optical profiling
    - Optical microscopy angle validation
    - Insert assembly
    - Bead sample preparation
    - Cell culture and seeding
    - Cell fixation
    - Fixed cell labeling for diffraction-limited imaging
    - Fixed cell labeling for DNA-PAINT imaging
    - Live cell labeling
    - Optical setup
    - Image acquisition settings
    - Image analysis and rendering
- Supplementary Figures
- Supplementary Tables
- Supplementary References

### SUPPLEMENTARY MATERIALS AND METHODS

**PDMS well insert fabrication.** SU8 molds with a ~330 μm height were prepared by spin-coating (WS-650Mz-23NPPB, Laurell Technologies) approximately 3 mL of the SU8-100 photoresist (Kayaku Advanced Materials Inc.) onto a silicon wafer first to 500 revolutions per minute (rpm) at 100 rpm/second acceleration, then held at 500 rpm for 10 seconds, then ramped to 1000 rpm at an acceleration of 300 rpm/second and held at this speed for 30 seconds. The molds were then heated on a hot plate at 65ºC for 4 hours followed by 95ºC overnight. A photomask (see Supplementary CAD files 6 and 7) was applied to the photoresist-coated silicon wafers which were then exposed to UV light under vacuum (140010, Vacuum UV-Exposure Box, Gie-Tec GmbH) with an 18 W light source for 45 seconds. The molds were then heated again on a hot plate at 65ºC for 1 minute followed by 95ºC for 20 minutes. The molds were then placed in a beaker with approximately 20 mL of SU8 developer (mr-Dev 600, Kayaku Advanced Materials, Inc.) and gently swirled for 25 minutes to dissolve residual untreated photoresist. All these steps were performed in accordance with the procedure outlined in the technical data sheet for SU8-100 negative epoxy photoresist (https://kayakuam.com/wp-content/uploads/2020/09/KAM-SU-8-50-100-Datasheet-9.3.20-Final.pdf), with minor deviations detailed above. After molds were completed, the height of the SU8 photoresist layer was determined using interferometric profilometry (NPFlex profiler, Bruker).

Prior to pouring the PDMS onto the SU8 molds, a macroscopic 3D-printed plastic cavity (see Supplementary CAD files 1 and 2) was added to enclose the mold, forming an insert with dimensions matching the imaging chamber. The cavity was fabricated using a Form2 printer (Formlabs) with Clear V4 resin (Formlabs), and a layer thickness of 0.025 mm was set in the PreForm software (Formlabs) for the print. The custom-designed piece laterally matched the SU8 photomask and axially matched the imaging chamber. For easy alignment, the cavity's bottom face was designed to feature grooves that matched the profiled height of the SU8 mold, allowing it to "click" into place atop the photoresist. This process created a PDMS pouring chamber that mimicked the imaging chamber dimensions while reserving an indent for micromirror placement. After the pieces were printed and washed with isopropyl alcohol (Form Wash, Formlabs) for 10 minutes, they were allowed to cure at 65ºC for 24 hours to prevent the printed material from inhibiting PDMS curing[1] before being placed on the SU8 mold.

PDMS (Sylgard 184 Silicone Elastomer Kit, Dow Inc.) was prepared at a 1:10 ratio of elastomer to curing agent and poured into the cavity where it was allowed to cure at 65ºC for 4 hours and carefully removed. A blade was used to carefully cut away any residual PDMS from the side of the insert that may have resulted from slight leakage from underneath the 3D-printed cavity.

**Micromirror fabrication.** Micromirror fabrication followed the same procedure described in Saliba & Gagliano *et al*[2]. Micromirror fabrication began with the design of structures of desired dimensions using AutoCAD (Autodesk, see Supplementary CAD files 3, 4, and 5). These designs were prepared for 3D nanoprinting in Describe (Nanoscribe GmbH & Co.KG) using the 10x Silicon Shell recipe using a slicing distance of 0.3 μm and a hatching distance of 0.3 μm. The dimensions of the micromirror were tailored to match those of the SU8 molds designed for each imaging well. The length and width of the micromirror were determined by the photomask dimensions, while the height was based on the profiling results of the SU8 photoresist mold using the NPFlex profiler (Bruker). This precise matching ensures a seamless fit of the micromirror within the PDMS well insert. For light sheet (LS) dimension characterization, micromirrors were designed with a sidewall angle of 45° and a height of 70 μm (for Ibidi chambers) or 100 μm (for

Mattek chambers), enabling a reflected LS with no tilt with respect to the image plane. For mammalian cell imaging, the micromirrors were fabricated with a 39° angle and 15 μm height, producing a reflected LS with a 12° tilt with respect to the image plane. Structures were fabricated via two-photon polymerization direct laser writing on a fused silica substrate, using the Nanoscribe Photonic Professional GT2 system and IP-Visio resin (Nanoscribe GmbH & Co.KG). Printing was performed with a 10x objective at 100% laser power and a scan speed of 40 mm/s. After printing, the micromirror structure was immersed briefly in SU8 developer to dissolve unpolymerized resin, followed by a rinse in isopropyl alcohol (Fisher Scientific) and drying with nitrogen gas. The micromirror structure was then treated for 15 minutes under an 18 W UV light source (Vacuum UV-Exposure Box, Gie-Tec GmbH) to cure any residual uncured resin. To prepare for vapor deposition, the structure was mounted on its side, ensuring only the edge containing the sidewall was metalized. This prevents unnecessary reflectivity from other surfaces. The structure was processed in an e-beam vapor deposition chamber, where 200 nm of silica (Kurt J. Lesker), 350 nm of aluminum (Kurt J. Lesker), and a final 5 nm layer of silica were deposited sequentially. The initial silica layer insulates the plastic insert, the aluminum provides reflectivity for single-objective LS reflection, and the final silica layer acts as a dielectric coating. This combination optimizes heat protection and enhances reflectivity.

**SEM imaging.** Scanning electron microscopy (SEM) (Apreo, ThermoFisher Scientific) was performed on inserts after metallization. Inserts were mounted on SEM chucks on a piece of metalized Kapton tape. To capture the morphology of the insert, the SEM stage was tilted at 25°. Secondary electron (SE) images were acquired at a 250x magnification using an Everhart-Thornley Detector (ETD), with an accelerating voltage (HV) of 5.00 kV and a probe current of 13 pA.

**Optical profiling.** Optical profilometry of the angled sidewalls of the micromirrors was performed using an NP Flex Optical Profilometer (Bruker) using a 5x objective. The sample was mounted on a custom 3D-printed angled wedge which has a flat surface of 8.5 mm*10 mm tilted at 45° (Form2 printer, Clear V4 resin, Formlabs), ensuring the profiling beam hit the mirror surface perpendicularly to accurately measure the intended face. A schematic of the mounting scheme can be found in Figure S3.

**Optical microscopy angle validation.** The angles of the inserts were validated using optical microscopy (Nikon Eclipse L150) with a 10X objective lens (Nikon LU Plan 10X). First, the insert was placed vertically on double-sided tape and imaged from the top to verify the reflective mirror angles. Line scans were made, and the angles between them were calculated in MATLAB to confirm agreement with the design specifications. Next, the insert was assembled with the PDMS and bonded to a glass coverslip (#1.5H, 22 × 22 $mm^2$, 170 ± 5 μm, 0102052, Marienfeld). The assembled insert was then taped onto a custom 3D-printed right-angle holder (Form2 printer, Clear V4 resin) and imaged from the side, allowing for visualization of the insert, the PDMS, and the glass surface (Supplementary Figure 4).

**Insert assembly.** To assemble the PDMS micromirror insert, the micromirror was placed in the correct orientation into the inset groove in the PDMS insert. Care must be taken to ensure the insert is seated in the correct orientation within the PDMS for LS reflection into the sample, which can be easily confirmed with a standard optical microscope. The assembled insert was secured in the imaging chamber through plasma bonding. For plasma bonding, the PDMS insert and glass-bottom imaging chamber were treated with air plasma (PDC-32G, Harrick Plasma Inc.) for one minute.

The insert was then aligned and pressed against the chamber bottom to form covalent bonds. These fabrication steps result in a robust and precisely aligned system, enabling effective LS imaging and cell culture within the imaging chamber.

**Bead sample preparation.** To compare the sectioning performance of epi-, HILO, and LS modalities, inserts featuring a 45° micromirror geometry were fabricated and assembled as described above. The imaging chambers were cleaned with argon plasma for 3 minutes and subsequently coated with a 0.02% gelatin solution prepared in mQ water (G1393, Sigma-Aldrich). The coating was allowed to air dry at 60°C to promote bead adhesion. Commercially available 15 µm surface-coated fluorescent microspheres (FocalCheck, F7235, Fisher Scientific) were diluted 1:10 in mQ water and allowed to settle in the gelatin-coated wells overnight. Prior to imaging, the solution was gently exchanged to PBS.

**Cell culture and seeding.** Following assembly, the imaging wells with inserts were sterilized by washing three times with 70% ethanol (BP82031GAL, Fisher Scientific) with a 5-minute incubation for each wash. This was followed by three washes with sterile PBS (SH3025601, Fisher Scientific) to remove the residual ethanol. The chambers were then coated with a 0.001% solution of fibronectin (F0895, Sigma Aldrich) in PBS by allowing the fibronectin solution to incubate in the chamber for 1 hour at 37°C. Human osteosarcoma cells (U-2 OS – HTB-96, ATCC) were then plated in imaging wells at a density of 100,000 cells/mL confirmed via cell counting with a hemocytometer (MDH-4N1, Sigma Aldrich). The cells were incubated at 37°C and 5% carbon dioxide (Thermo Scientific Heracell 150i $CO_2$ Incubator, 51-032-871, Fisher Scientific) in high-glucose Dulbecco's modified Eagle's medium (DMEM, Gibco) with 25 mM HEPES and supplemented with 10% (v/v) fetal bovine serum (FBS, Gibco) and 1 mM sodium pyruvate (Gibco).

**Cell fixation.** After incubation for 12 hours, the cells were washed three times with sterile PBS and fixed for 15 minutes at 37°C and 5% carbon dioxide with 4% formaldehyde solution made by dissolving paraformaldehyde (16% PFA, Electron Microscopy Sciences) in PBS. After fixation, the fixed cells were washed three times with sterile PBS and treated with 10 mM ammonium chloride (Sigma Aldrich) in PBS for 10 minutes. The fixed cells were then stored in PBS at 4°C until labeling and imaging.

**Fixed cell labeling for diffraction-limited imaging.** Cells were immunolabeled by first being permeabilized with 0.1% saponin (SAE0073, Sigma Aldrich) in PBS for 10 minutes. For diffraction-limited lamin B1 imaging, cells were then blocked with 3% (w/v) bovine serum albumin (BSA) (A2058-5G, Sigma Aldrich) and 10% donkey serum (ab7475, abcam) in 0.1% saponin in PBS. Cells were then labeled with rabbit anti-lamin B1 (ab16048, Abcam) primary antibodies using a 1:1,000 dilution in 1% (w/v) BSA and 10% donkey serum in 0.1% saponin for 3 hours. Cells were then washed five times with PBS and labeled with donkey anti-rabbit secondary antibodies conjugated with dye CF568 (20098-1, Biotium) at a 1:100 dilution in PBS for 1 hour. Finally, cells were washed ten times with PBS to remove excess secondary antibodies.

**Fixed cell labeling for DNA-PAINT imaging.** For single-molecule DNA-PAINT imaging, after permeabilization with 0.1% saponin in PBS for 10 minutes, cells were blocked with a DNA-PAINT blocking buffer comprising 3% (w/v) BSA (A2058-5G, Sigma Aldrich), 0.05 mg/mL sheared salmon sperm DNA (15632011, Invitrogen, ThermoFisher Scientific), 0.02% Tween-20 (P1379-500ML, Sigma Aldrich), and 0.05% sodium azide (50-103-7106, G-Biosciences, Fisher Scientific) in 0.1% saponin for 1 hour.

Subsequent labeling was performed for each target using a primary antibody that was preincubated with a secondary nanobody conjugated to a DNA oligonucleotide docking strand. Nanobodies conjugated to docking strands were prepared using the procedures outlined in Sograte-Idrissi *et al*[3]. and Schlichthaerle *et al*[4]. In brief, anti-rabbit nanobodies with an unconjugated C-terminal cysteine (NanoTag) were diluted in 5 mM Tris(2-carboxyethyl)phosphine hydrochloride (TCEP) (C4706-2G, Sigma Aldrich). The nanobodies were then filtered from the solution via centrifugation using Amicon spin filters (10 kDa MWCO, UFC501024, Sigma Aldrich) and incubated with a 50 molar excess of dibenzocyclooctyne (DBCO)-maleimide crosslinker (760668-5MG, Sigma Aldrich) overnight at 4°C. The nanobodies were then filtered from solution via centrifugation using Amicon spin filters and incubated with a 10 molar excess of azide-DNA oligonucleotide strands (Integrated DNA Technologies) for 2 hours at room temperature. The final nanobodies conjugated to oligonucleotide docking strands were exchanged into PBS using centrifugation with Amicon spin filters.

For TOMM20 imaging, rabbit anti-TOMM20 primary antibodies diluted at 1:200 (ab186735, Abcam) were premixed with anti-rabbit secondary nanobodies conjugated to docking strands at a 0.01 μg/μL concentration in labeling buffer (1% (w/v) BSA, 0.05 mg/mL sheared salmon sperm DNA, and 0.05% sodium azide in 0.1% saponin) for 1 hour. The premixed solution was then added to the cells after blocking and allowed to incubate for 3 hours. The cells were then washed ten times with PBS prior to imaging.

For two-target TOMM20 and vimentin imaging, rabbit anti-TOMM20 primary antibodies diluted at 1:200 (ab186735, Abcam) and rabbit anti-vimentin primary antibodies (ab92547, abcam) diluted at 1:100 were premixed separately with anti-rabbit secondary nanobodies conjugated to orthogonal docking strands for each respective target at a 0.01 μg/μL concentration in labeling buffer (1% (w/v) BSA, 0.05 mg/mL sheared salmon sperm DNA, and 0.05% sodium azide in 0.1% saponin) for 1 hour. The premixed TOMM20 solution was then added to the cells after blocking and allowed to incubate for 3 hours. The cells were then washed ten times with PBS. The premixed vimentin solution was then added to the cells and allowed to incubate for 3 hours. The cells were then washed ten times with PBS prior to imaging.

For DNA-PAINT acquisitions, after labeling and prior to imaging, 0.1 μm 580/605 nm fiducial beads (F8801, Invitrogen) were added at a 1:1,000,000 dilution in PBS and allowed to settle for 5 minutes to enable adhesion to the coverslip for drift correction. The bead solution was then aspired from the sample, followed by three washes with PBS to remove unadhered beads.

**Live cell labeling.** After incubation for 12 hours, the cells were washed with warm cell media and incubated with a 200 nM solution of MitoTracker Orange CMTMRos (M7510, ThermoFisher) in cell media for 30 minutes at 37°C and 5% carbon dioxide. The cells were then washed three times with warm cell media to remove excess MitoTracker dye prior to imaging.

**Optical setup.** The overall single-objective LS optical setup design used is described in detail in Saliba & Gagliano *et al*[2]. Here, the optical platform was built around a conventional inverted microscope (Ti2-E Inverted Microscope, Nikon) using a 100x, NA 1.49 objective (MRD01995 Apo TIRF, Nikon) with a custom-built excitation and emission pathway. In brief, a Gaussian LS, whose dimensions are tuned using lenses in the optical path, was formed using a cylindrical lens using a 560 nm laser (560 nm, 1000 mW, MPB Communications). It was steered with two galvanometric mirrors and a tunable lens conjugated to the back focal plane of the objective, enabling easy alignment of the reflected LS in the sample. An additional galvanometric mirror conjugated to the sample plane was used to dither the LS at a frequency of 100 Hz. Flip mirrors in the excitation path were used to easily switch between LS and epi-illumination modalities. A translation stage (LT1, Thorlabs) was used to translate the flip mirror and lens prior to the objective in the epi- path to achieve HILO illumination. In the detection path, a tube lens, followed by a 4f lens system, focused emitted light onto a sCMOS camera (Prime95B, Teledyne Photometrics). The effective pixel size of our system was calibrated by imaging a 600 line pairs per mm Ronchi ruling (38-566 600 lp/mm, Edmund Optics) using the camera. This resulted in a calibrated pixel size of 110 nm/pixel. The average conversion gain of the camera was calculated to be 0.54 photoelectrons per analog-to-digital (A/D) count, and the base level was calculated based on experimental data as 80 A/D counts. Two-target single-molecule imaging was performed using the same optical system, but with an EMCCD camera (iXon Ultra 897, Andor, Oxford Instruments) with a set EM gain of 200, which corresponded to a calibrated EM gain of 182. The conversion gain was determined to be 4.41 photoelectrons per A/D count, the base level was calculated as 473 A/D counts, and the calibrated pixel size was 159 nm/pixel. To image the LS in fluorescent solution, images were acquired using a CMOS camera (CS235MU, Thorlabs) with a calibrated pixel size of 59 nm/pixel operated with the ThorCam software (version 3.7.0.6).

**Image acquisition settings.** For all data sets, except for imaging of the LS in a fluorescent solution, where the ThorCam software (version 3.7.0.6) was used, Micromanager[5,6] was used to control camera settings and exposure time. For imaging of 15 µm fluorescent beads, beads were imaged with a 560 nm laser at ~40 mW/cm$^2$ and an exposure time of 100 ms. For diffraction-limited imaging of lamin B1 in U2OS cells, cells were imaged with a 560 nm laser at ~1.8 W/cm$^2$ and an exposure time of 100 ms.

For TOMM20 DNA-PAINT epi- versus LS illumination imaging, the 560 nm laser was used at ~50 W/cm$^2$ for epi-illumination and LS illumination using an exposure time of 150 ms to roughly match the binding kinetics of the transiently binding imager strands. 10,000 frames were acquired for both epi- and LS illumination.

For two-target TOMM20 and vimentin imaging, the 560 nm laser was used at ~60 W/cm$^2$ with LS illumination using an exposure time of 200 ms. 20,000 frames were acquired for each target. Vimentin was imaged first followed by TOMM20. Imager strands were manually removed via micropipetting on the microscope stage. To ensure minimal crosstalk between targets, 15 extensive washes were performed with imaging buffer (PBS with 500 mM NaCl, pH 8.0) before adding strands for the next target. Crosstalk analysis was performed on an 18 µm x 20 µm area across 500 frames for vimentin, the wash, and TOMM20. The resulting localization counts were 12,833, 278, and 5,763, respectively, confirming the efficacy of the washing protocol in removing residual imager strands between targets.

Imager strands for all DNA-PAINT acquisitions were custom-ordered conjugated to the dye Cy3B (Integrated DNA Technologies). Sequences can be found in Supplementary Table 2. For all DNA-PAINT acquisitions, imager strands were prepared at a concentration of 0.1 nM in imaging buffer (PBS with 500 mM NaCl, pH 8.0) supplemented with Trolox[7]. Trolox was prepared at a 100x stock concentration and aliquoted and stored at -20°C for up to 6 months using the procedure outlined by Steen *et al*[8]. Aliquots were thawed and diluted 1:100 in imaging buffer and the solution was allowed to rest for at least 15 minutes prior to introducing imager strands and imaging.

For live cell imaging, cells were imaged with a 560 nm laser at ~0.1 W/cm$^2$ using LS illumination in 30 second intervals using 100 frames at an exposure time of 50 ms for every interval for 10 minutes.

**Image analysis and rendering.** All image data were processed using the open-source software ImageJ[9,10]. Single-molecule DNA-PAINT data were analyzed in ThunderSTORM[11], which is an open-source ImageJ plugin, using wavelet filtering for background subtraction and a weighted least-squares fitting routine. PSF detection was performed with a local maximum fitting method with a peak intensity threshold coefficient of 2.2. For drift correction, the tracked motion of the fiducial bead was smoothed with a cubic spline fitting function and subtracted from localized single-molecule data. The custom written codes for drift correction are available on GitHub (https://github.com/Gustavsson-Lab/2D-Drift-Correction). For two-target data, the position of the bead was also used to correct for drift between targets. For epi- versus LS comparison, single-molecule data were filtered in ThunderSTORM for intensity greater than 100 and less than 5000 photons per localization. For two-target imaging, single-molecule data were filtered in ThunderSTORM for intensity greater than 400 and less than 6000 photons per localization and localization precision, or uncertainty, between 5 and 25 nm to remove bad localizations. The data sets were also filtered for 10-nearest neighbors with a denoise range of 0.01-20 to remove spurious localizations. Fourier ring correlation (FRC) analysis was performed in Vutara SRX (Bruker) with a pixel size of 8 nm and a threshold of 0.143 (1/7). Two-target data were rendered in Vutara SRX with a particle size of 50 nm and an opacity of 0.65 for TOMM20 and an opacity of 0.45 for vimentin. All other datasets were rendered in ImageJ using the mpl-inferno look up table.

### SUPPLEMENTARY FIGURES

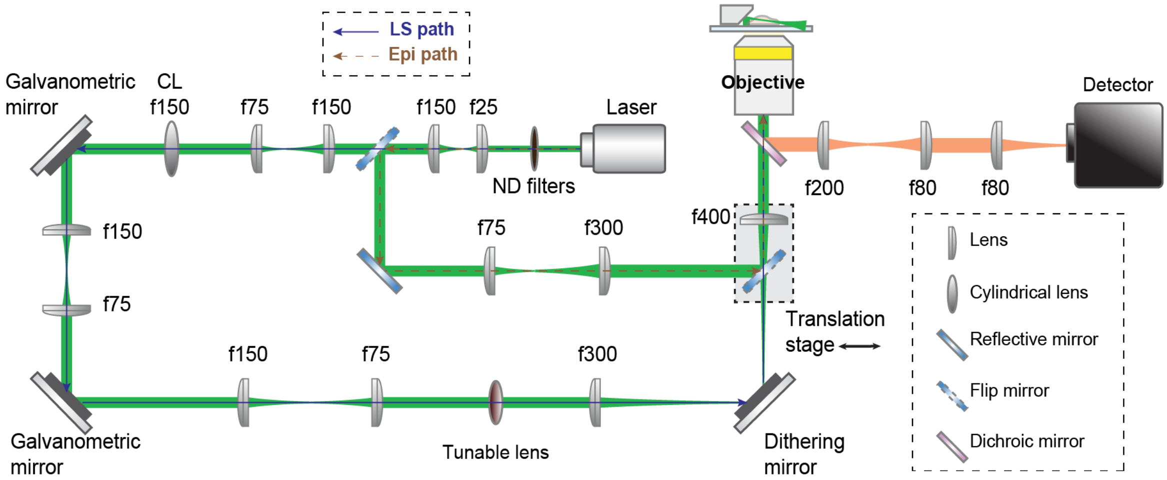


**Supplementary Figure 1.** Schematic of the optical setup used for single-objective light sheet (LS), highly inclined and laminated optical sheet (HILO), and epi-illumination imaging. In the epi- and HILO illumination path, a translation stage is used to switch between epi- and HILO illumination modalities. In the LS illumination path, the LS is formed with a cylindrical lens (CL) and reflected by galvanometric mirrors conjugated to the back focal plane of the objective lens for steering. The focus of the beam is adjusted using the tunable lens, which is also conjugated to the back focal plane of the objective lens. The LS is dithered by a galvanometric dithering mirror conjugated to the sample plane. Switching between LS and epi-illumination pathways is enabled by flip mirrors. Emitted light is collected by the objective lens in the microscope, focused by a tube lens (f200), and passed through a 4*f* system before being imaged on an sCMOS or EMCCD camera.

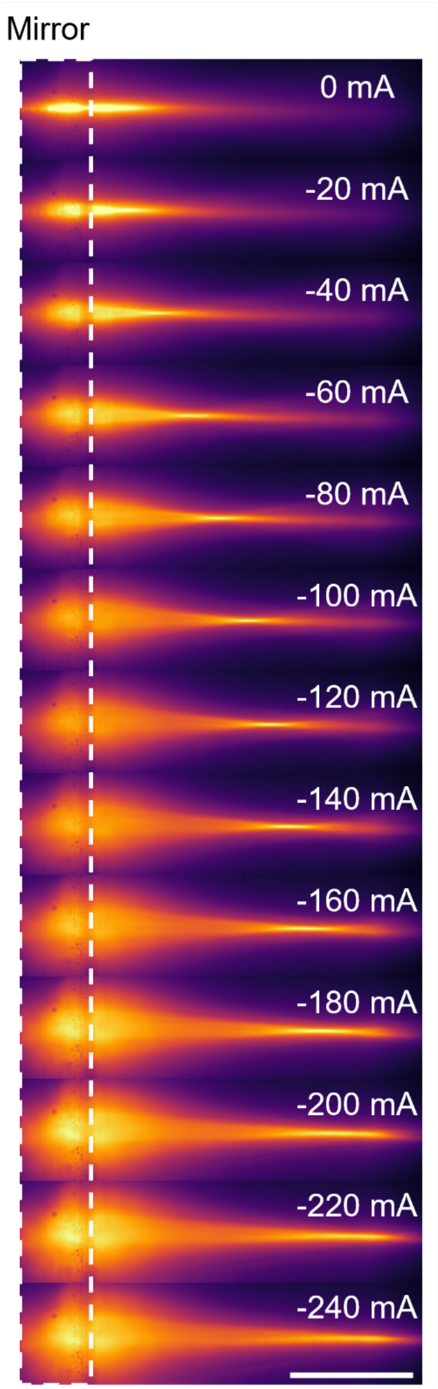


**Supplementary Figure 2.** Characterization of the tunability of the position of the light sheet (LS) beam waist. Images demonstrating the displacement of the LS beam waist as a function of the tunable lens current. By modulating the current from 0 mA to -240 mA in 20 mA increments, the focal position of the beam waist is translated relative to the micromirror surface, providing a robust range up to ~50 µm for sample-specific alignment. Scale bar is 20 µm.

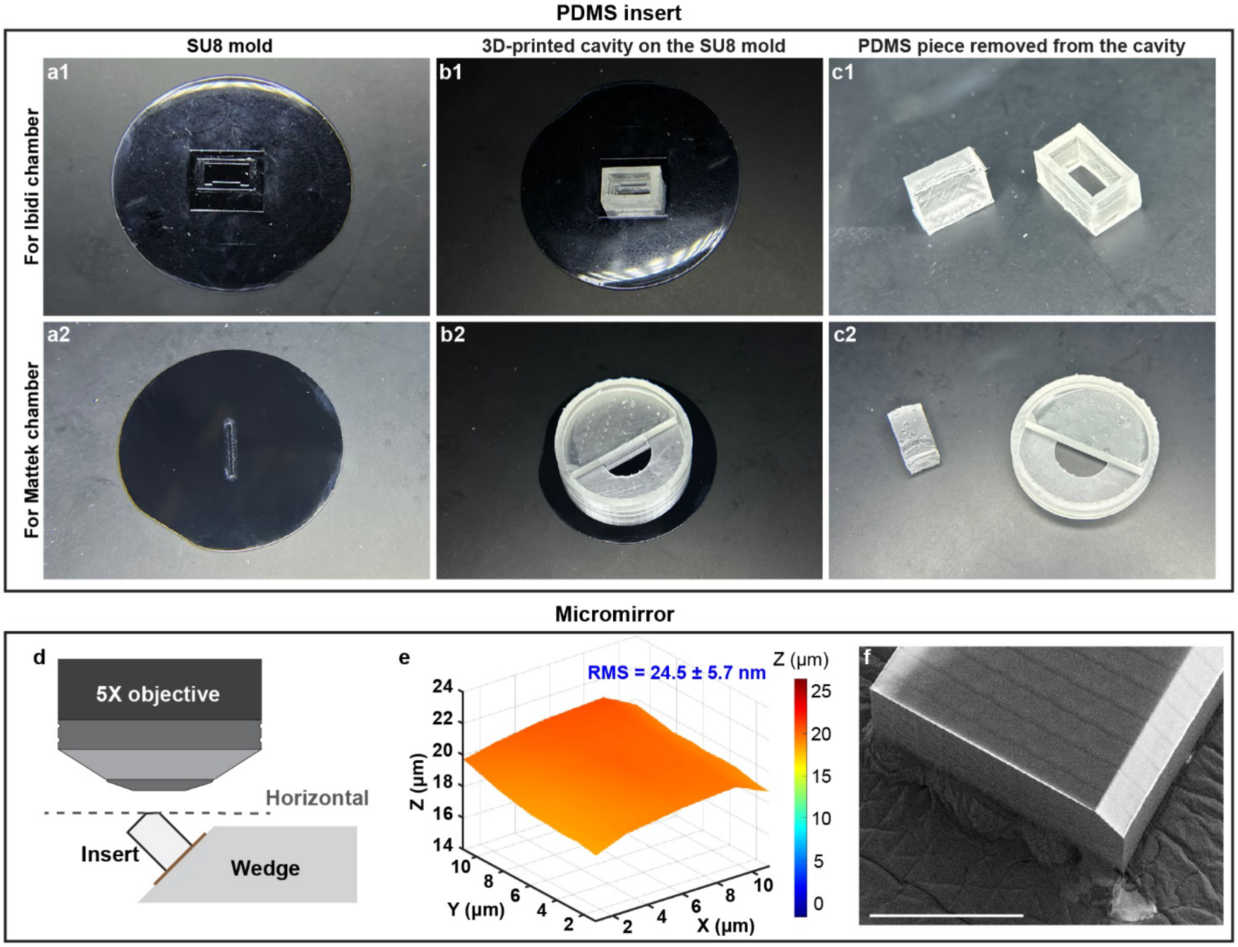


**Supplementary Figure 3.** Details of the fabrication procedure and characterization of micromirror geometry and roughness. (a1-a2) Photos of the SU8 molds made on Si wafers. (b1-b2) 3D-printed cavities and SU8 molds are assembled to make the PDMS pieces. (c1-c2) PDMS pieces are removed from the 3D-printed cavities, and extra parts are cut out. (d) Schematic of the optical profilometry approach used to measure the 3D profile of the metalized micromirror surface and (e) characterization of the surface roughness. (f) SEM image of the micromirror insert. Scale bar is 300 μm.

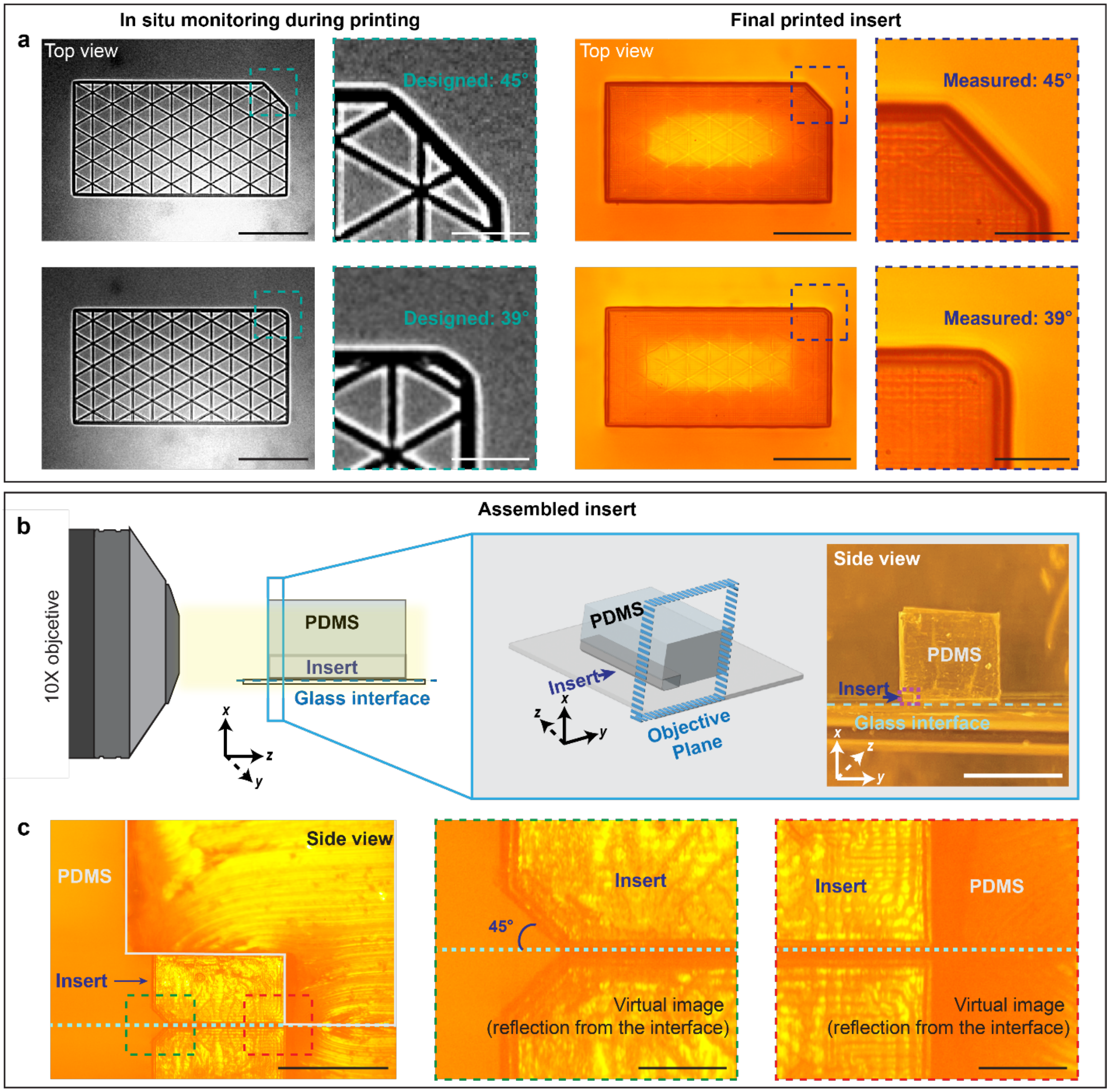


**Supplementary Figure 4.** Experimental validation of micromirror geometry. (a) Images captured during (left) and after (right) the 3D nanoprinting process, confirming precise angular geometries for both 45° (top) and 39° (bottom) micromirror designs. Scale bars are 200 μm for full images and 50 μm for insets. (b) Schematics and image of the optical microscopy configuration used to characterize the micromirror angle after bonding to the coverslip. Scale bar is 5 mm. (c) Optical microscopy images validating the flush interface between the micromirror and the coverslip within the PDMS insert, confirming the intended 45° reflection geometry. Scale bars are 500 μm (left) and 100 μm (center/right).

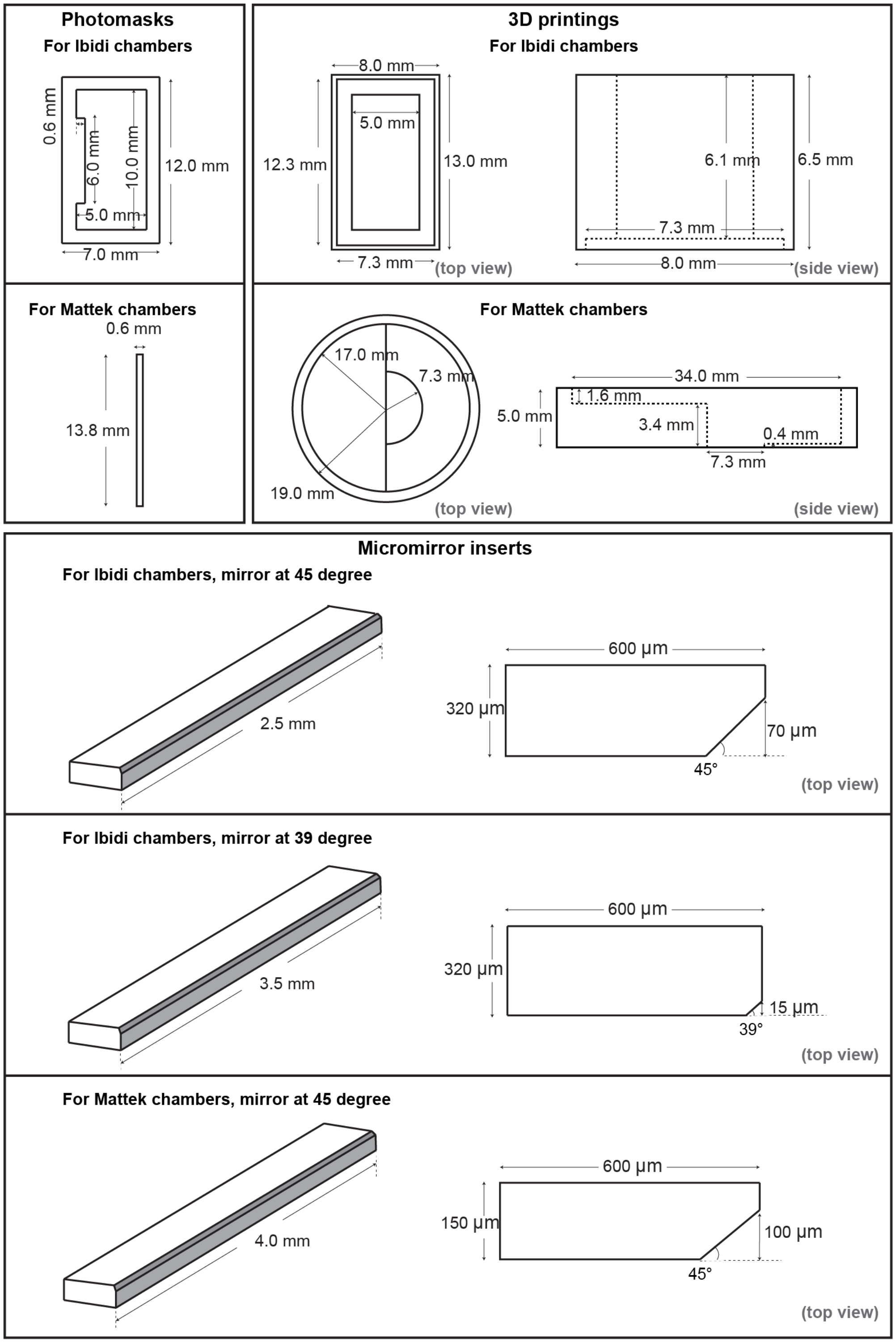


**Supplementary Figure 5.** Detailed schematics with labeled dimensions of each of the components for both the Ibidi and Mattek insert fabrication pipelines. The drawings are not to scale.

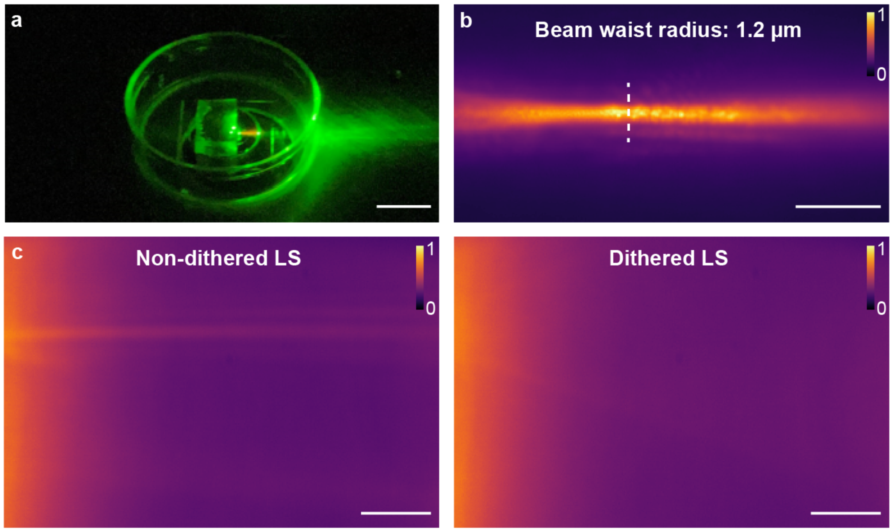


**Supplementary Figure 6.** Demonstration of inserts enabling single-objective light sheet (LS) reflection in a circular Mattek chamber. (a) Photo of the Mattek chamber with an insert on a microscope stage reflecting a single-objective LS in fluorescent solution. Scale bar is 10 mm. (b) The thin end of the LS imaged in a fluorescent solution using the Mattek chamber. (c) The wide end of the LS used for optical sectioning imaged in a fluorescent solution using the Mattek chamber without versus with dithering, revealing the homogenization of the LS intensity upon dithering. Scale bars in (b) and (c) are 10 μm. The colorbars show intensity normalized independently for each image.

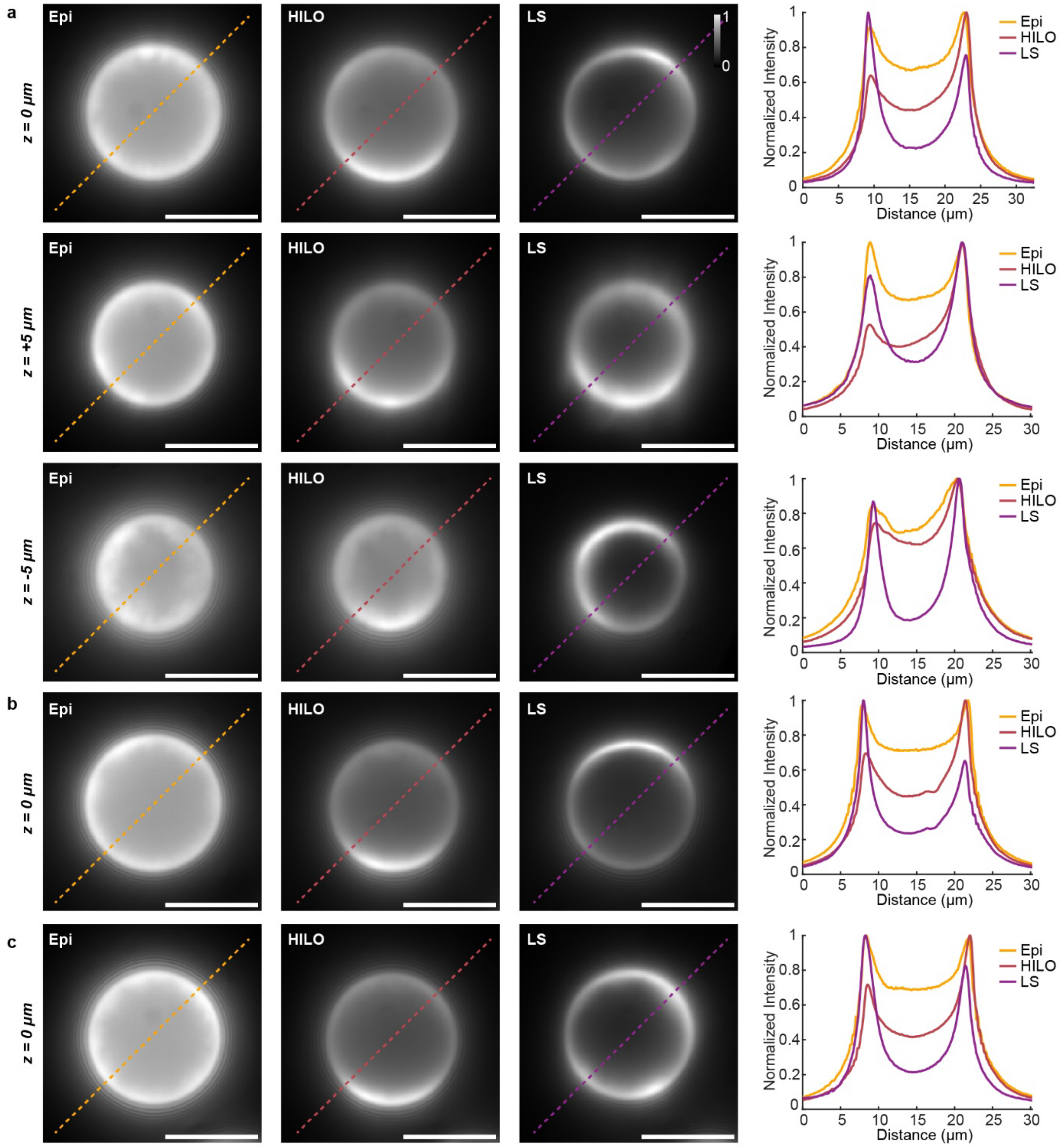


**Supplementary Figure 7.** Comparison of signal-to-background ratio (SBR) across different illumination modalities. (a) Images of a 15 µm surface-labeled fluorescent bead under epi-, highly inclined and laminated optical sheet (HILO), and light sheet (LS) illumination. SBR is compared across three axial planes ($z$ = 0, +5, and -5 µm, where 0 µm represents the center of the bead). The LS modality maintains superior sectioning and SBR across all planes, whereas HILO performance degrades at defocused positions. (b-c) Technical replicates of additional beads at the $z$ = 0 µm plane, confirming the reproducibility of the observed performance trends. Scale bars are 10 µm.

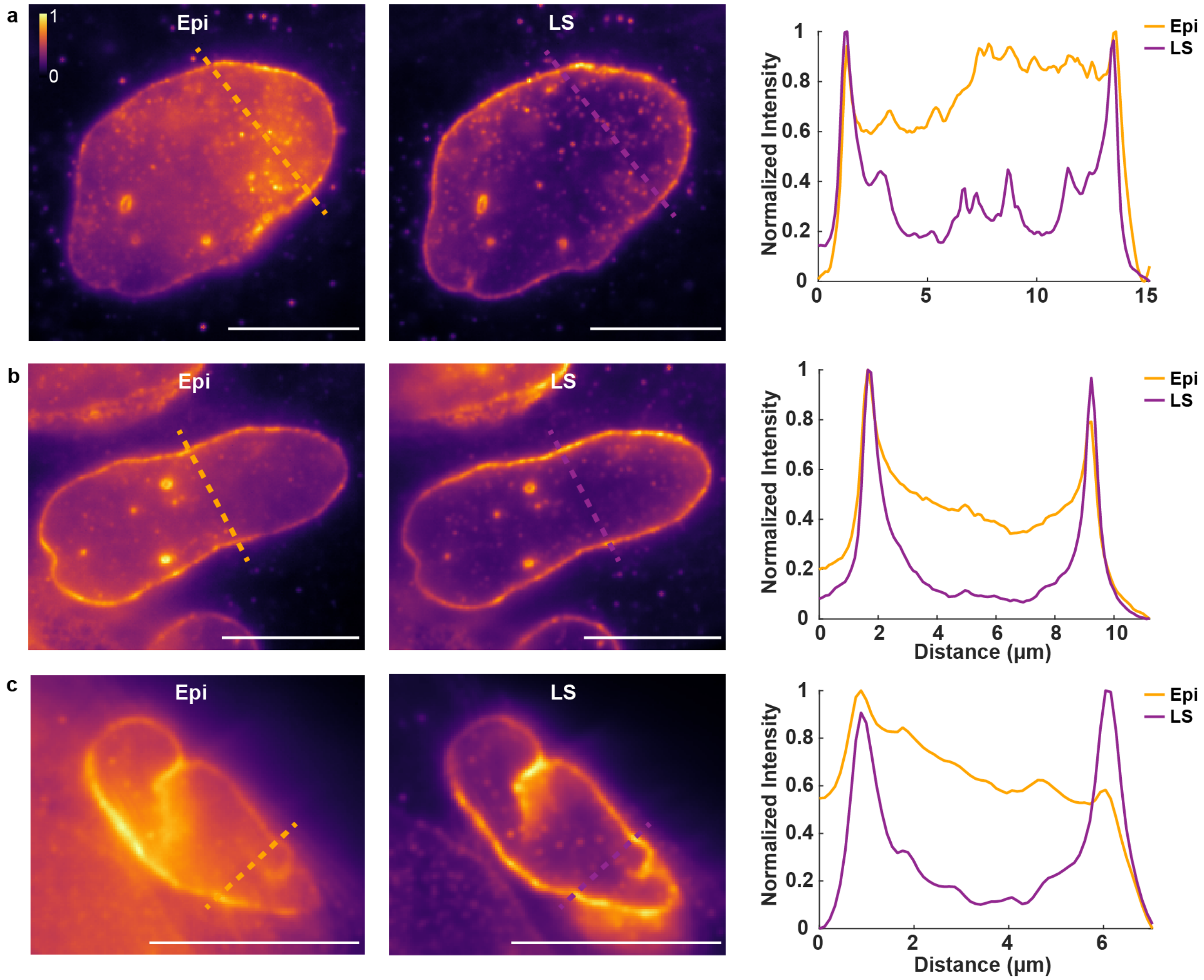


**Supplementary Figure 8.** Diffraction-limited images with line scans of lamin B1 imaged with single-objective light sheet (LS) or epi-illumination. (a-c) Images with line scans and corresponding graphs of the normalized intensity demonstrate consistently improved signal-to-background ratio (SBR) with LS compared with epi-illumination, as shown in Figure 2b. Scale bars are 10 µm. The colorbar shows intensity normalized independently for each image.

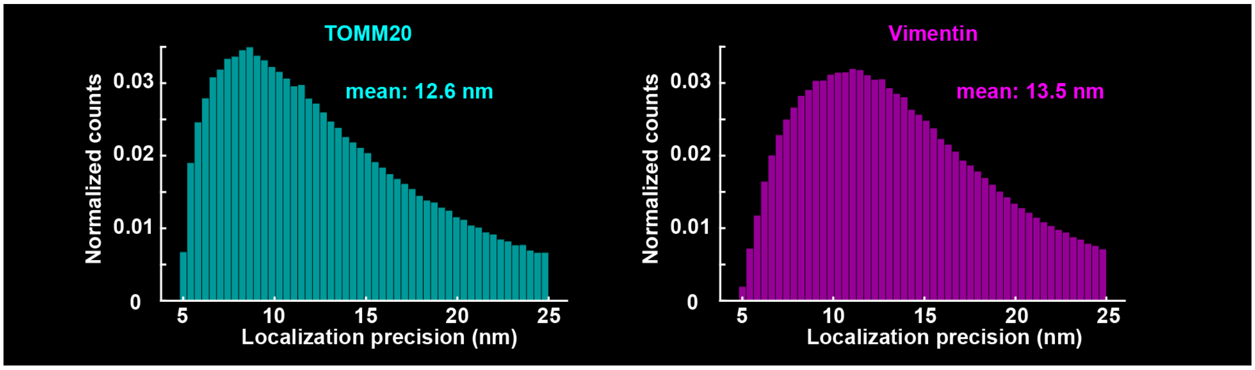


**Supplementary Figure 9.** Localization precision histograms for two-target single-molecule super-resolution imaging of TOMM20 and vimentin shown in Figure 3. The mean values of the distributions are shown for each histogram.

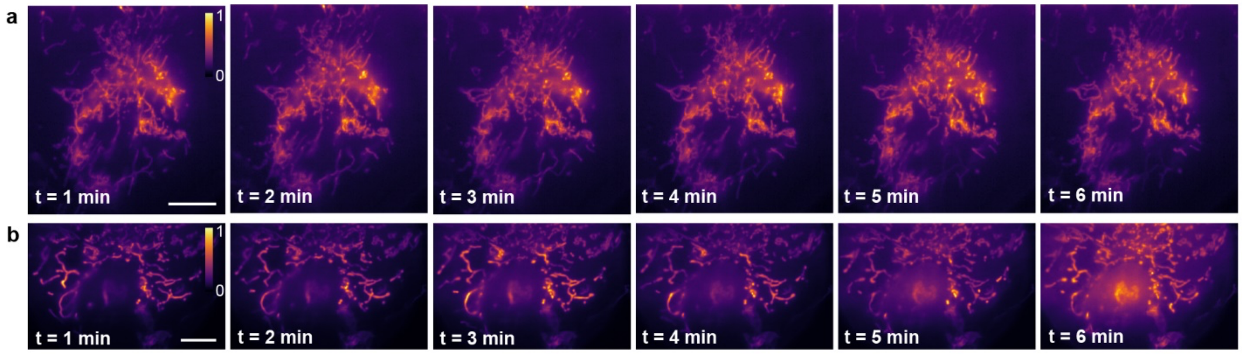


**Supplementary Figure 10.** Time lapse images of mitochondria imaged with a single-objective light sheet (LS) in two separate live cells in (a) and (b). Scale bars are 10 μm. The colorbars show intensity normalized independently for each image.

### SUPPLEMENTARY TABLES

**Supplementary Table 1.** Parameters used to prepare micromirrors for 3D printing with 2-photon polymerization. Micromirror structures were prepared in Describe prior to printing using the IP-Q 10x Silicon Shell (3D LF) recipe. Splitting mode was set to none and the laser power and scan speed were set to be the same for shell, scaffold, and base writing.

| Slicing distance | Hatching | Base slice count | Shell contour count | Walls scaffolding (nm) | Floors scaffolding (nm) | Laser power (%) | Scan speed (mm/s) |
|---|---|---|---|---|---|---|---|
| 0.3 µm | Distance: 0.3 µm Angle: 45° Angle Offset: 0° | 5 | 20 | Spacing: 55 Thickness: 5 | Spacing: 60 Thickness: 5 | 100 | 40,000 |

**Supplementary Table 2.** Docking and imager sequences used for single-molecule DNA-PAINT imaging.

| Target imaged | Docking sequence | Imager sequence |
|---|---|---|
| TOMM20 | nanobody-TTATACATCTA-3' | 5'-CTAGATGTAT-dye |
| Vimentin | nanobody-TTTCTTCATTA-3' | 5'-GTAATGAAGA-dye |

### SUPPLEMENTARY REFERENCES


(1) Venzac, B.; Deng, S.; Mahmoud, Z.; Lenferink, A.; Costa, A.; Bray, F.; Otto, C.; Rolando, C.; Le Gac, S. PDMS Curing Inhibition on 3D-Printed Molds: Why? Also, How to Avoid It? *Anal. Chem.* **2021**, *93* (19), 7180–7187. https://doi.org/10.1021/acs.analchem.0c04944.

(2) Saliba, N.; Gagliano, G.; Gustavsson, A.-K. Whole-Cell Multi-Target Single-Molecule Super-Resolution Imaging in 3D with Microfluidics and a Single-Objective Tilted Light Sheet. *accepted, Nat. Commun.* **2024**. https://doi.org/10.1101/2023.09.27.559876.

(3) Sograte-Idrissi, S.; Oleksiievets, N.; Isbaner, S.; Eggert-Martinez, M.; Enderlein, J.; Tsukanov, R.; Opazo, F. Nanobody Detection of Standard Fluorescent Proteins Enables Multi-Target DNA-PAINT with High Resolution and Minimal Displacement Errors. *Cells* **2019**, *8* (1), 48. https://doi.org/10.3390/cellold010048.

(4) Schlichthaerle, T.; Strauss, M. T.; Schueder, F.; Auer, A.; Nijmeijer, B.; Kueblbeck, M.; Jimenez Sabinina, V.; Thevathasan, J. V.; Ries, J.; Ellenberg, J.; Jungmann, R. Direct Visualization of Single Nuclear Pore Complex Proteins Using Genetically-Encoded Probes for DNA-PAINT. *Angew Chem Int Ed* **2019**, *58* (37), 13004–13008. https://doi.org/10.1002/anie.201905685.

(5) Edelstein, A.; Amodaj, N.; Hoover, K.; Vale, R.; Stuurman, N. Computer Control of Microscopes Using µManager. *CP Molecular Biology* **2010**, *92* (1). https://doi.org/10.1002/0471142727.mb1420s92.

(6) Edelstein, A. D.; Tsuchida, M. A.; Amodaj, N.; Pinkard, H.; Vale, R. D.; Stuurman, N. Advanced Methods of Microscope Control Using µManager Software. *J Biol Methods* **2014**, *1* (2), e10. https://doi.org/10.14440/jbm.2014.36.

(7) Cordes, T.; Vogelsang, J.; Tinnefeld, P. On the Mechanism of Trolox as Antiblinking and Antibleaching Reagent. *J. Am. Chem. Soc.* **2009**, *131* (14), 5018–5019. https://doi.org/10.1021/ja809117z.

(8) Steen, P. R.; Unterauer, E. M.; Masullo, L. A.; Kwon, J.; Perovic, A.; Jevdokimenko, K.; Opazo, F.; Fornasiero, E. F.; Jungmann, R. The DNA-PAINT Palette: A Comprehensive Performance Analysis of Fluorescent Dyes. *Nat Methods* **2024**, *21* (9), 1755–1762. https://doi.org/10.1038/s41592-024-02374-8.

(9) Abramoff, M.; Magalhães, P.; Ram, S. J. Image Processing with ImageJ. *Biophotonics International* **2003**, *11*, 36–42.

(10) Schneider, C. A.; Rasband, W. S.; Eliceiri, K. W. NIH Image to ImageJ: 25 Years of Image Analysis. *Nat Methods* **2012**, *9* (7), 671–675. https://doi.org/10.1038/nmeth.2089.

(11) Ovesný, M.; Křížek, P.; Borkovec, J.; Švindrych, Z.; Hagen, G. M. ThunderSTORM: A Comprehensive ImageJ Plug-in for PALM and STORM Data Analysis and Super-Resolution Imaging. *Bioinformatics* **2014**, *30* (16), 2389–2390. https://doi.org/10.1093/bioinformatics/btu202.
